# Sequence based prediction of protein phase separation into disordered condensates using machine learning

**DOI:** 10.1101/2021.12.13.472521

**Authors:** Pratik Mullick, Antonio Trovato

**Affiliations:** Department of Physics and Astronomy ‘Galileo Galilei’, University of Padova, Padova PD, Italy

## Abstract

Several proteins which are responsible for neuro-degenrerative disorders (Alzheimer’s, Parkinson’s etc) are shown to undergo a mechanism known as liquid liquid phase separation (LLPS). We in this research build a predictor which would answer whether a protein molecule would undergo LLPS or not. For this we used some protein sequences for which we already knew the answer. The ones who undergo LLPS were considered as the positive set and the ones who do not, were taken as the negative set. Depending on the knowledge of amino-acid sequences we identified some relevant variables in the context of LLPS e.g. number of amino acids, length of the best pairings, average register shifts. Using these variables we built a number of scoring functions which were basically analytic functions involving these variables and we also combined some scores already existing in the literature. We considered a total of 43636 protein sequences, among them only 121 were positive. We applied logistic regression and performed cross validation, where 25% of the data were used as the training set and the performance of the obtained results were tested on the remaining 75% of the data. In the training process, we used Simplex algorithm to maximize area under the curve (AUC) in receiver operator characteristics (ROC) space for each of the scores we defined. The optimised parameters were then used to evaluate AUC on the test set to check the accuracy. The best performing score was identified as the predicting model to answer the question whether a protein chain would undergo phase separating behavior or not.

**Author summary:** Liquid-liquid phase separation (LLPS) is a classic subject in polymer physics. The essential physics is aptly captured within the Flory-Huggins (FH) approach, a simple lattice mean-field theory. Evidence has been mounting in the last decade that protein LLPS underlies the formation of membrane-less organelles (MLOs) in living cells, explaining why proteins and other bio-molecules can remain in a dense liquid condensate without diffusing away. Intrinsically disordered regions (IDRs), with a sequence-intrinsic preference for conformational heterogeneity or disorder under native conditions, are enriched in proteins that have the ability of switching on LLPS. The detailed understanding of the biological function of disordered bio-molecular condensates, whose formation is driven by LLPS, is currently the focus of a major effort in cell biology. Several key proteins in neuro-degenerative disorders are components of MLOs, and a further liquid-to-solid transition to unsoluble amyloid aggregates may be triggered by pathogenic mutants. Non-equilibrium active processes are also known to drive non trivial spatio-temporal organization patterns in MLOs. In this research we aim to predict which proteins can undergo LLPS in physiological conditions in living cells, and the corresponding phase behavior, based on theoretical tools and on the knowledge of the amino-acid sequence alone. We derive specific knowledge-based potentials for the different kind of short-range interactions that are believed to drive protein LLPS and use them to build a simple yes/no predictor of phase separation in physiological conditions.

## Introduction

Liquid-liquid phase separation (LLPS), or demixing, is a classic subject in polymer physics [1]. The essential physics is aptly captured within the Flory-Huggins (FH) approach, a simple lattice theory, where the free energy of mixing per lattice site can be derived using a mean-field assumption [2, 3]. The driving force underlying LLPS is the exchange of chain/solvent interactions for chain/chain and solvent/solvent interactions under conditions for which this process is energetically favorable, a balance quantified by the Flory parameter.

Evidence has been mounting in the last decade that protein LLPS underlies the formation of membrane-less organelles (MLOs) in living cells [4]. In fact, eukaryotic cells are composed of numerous compartments or organelles. These organelles carry out specific functions and provide spatio-temporal control over cellular materials, metabolic processes, and signaling pathways. For example, the nucleus physically separates DNA transcription from RNA translation; other examples of membrane-bound organelles include lysosomes, the endoplasmic reticulum, and synaptic vesicles. However, cells also harbor several NLOs that lack a delimiting membrane. These are supra-molecular assemblies composed of proteins, nucleic acids, and other molecular components, that are present in the nucleus as well as in the cytoplasm. One feature that has attracted considerable attention is the presence of intrinsically disordered regions (IDRs) in proteins that have the ability of switching on LLPS [5]. These regions display a sequence-intrinsic preference for conformational heterogeneity or disorder under native conditions [6]. The detailed understanding of the biological function of disordered bio-molecular condensates, whose formation is driven by LLPS, is currently the focus of a major effort undertaken by a large community in cell biology [7].

In particular, several key proteins in neuro-degenerative disorders are components of MLOs [8]. The observed conversion of dynamic protein droplets to solid aggregates [9] shows them to be meta-stable or inherently unstable, and shows that specific cellular processes keep them from solidifying. These liquid-to-solid transitions are accelerated by disease mutations [10] that seem to target *β*-zippers in IDRs [11], which makes them more prone to fold into stable amyloid structures [12]. Amyloid fibrils contain pairs of closely mating *β*-sheets along the fibril axis, and their presence is implicated in several degenerative pathologies triggered by aberrant protein mis-folding and subsequent aggregation [13].

There is much debate on the type of interactions that underlie protein LLPS at a molecular level. Multi-valent interactions of heterogeneous modular binding domains and their target motifs can drive LLPS of proteins with a well-defined native structure [14], yet the forces promoting LLPS of IDRs are less understood. A role has been suggested for several weak non-covalent interactions such as electrostatic, dipole-dipole, pi-pi stacking, cation-pi, hydrophobic and hydrogen bonding (namely *β*-zipper) interactions [4]. In particular, the importance of charge patterns has been highlighted [15].

In a broader perspective, and at a larger scale, a possible role has been hypothesized for non-equilibrium active processes in MLOs, to prevent pathological fibril formation and gelation in MLOs [5]. Moreover, ongoing chemical reactions that are catalyzed by enzymes in the cell can also alter the equilibrium picture, whereby a system containing several droplets will generally evolve by larger droplets growing at the expense of smaller ones, a process termed Ostwald ripening [16]. As molecules are constantly exchanging between droplets and solution, ripening occurs by a net diffusion of molecules out of smaller droplets and into more stable larger ones. The influence of Ostwald ripening can be opposed by chemical reactions that produce or deplete droplet material. For example, a FH approach that included chemical reactions was used to model the dynamics of centrosomes, which are important in organizing networks of microtubules in cells [17].

A central issue in the field is the ability of predicting which proteins can undergo LLPS in physiological conditions in living cells, based on the knowledge of the amino-acid sequence alone, in particular for IDRs [7]. The understanding of the sequence determinants of phase separation in IDRs is still rudimentary, but it is clear that different flavors of IDRs exist that determine the type of stimulus the IDR responds to [18]. The sequence also likely determines the emergent properties of its dense phase, i.e., dense-phase concentration [19], and material properties such as visco-elasticity [20]. In this respect, theoretical advances have been somewhat limited, despite the problem being tackled with several approaches at different coarse-graining levels. Langevin molecular dynamics simulations, coupled with enhanced sampling methods, have been used for the direct extraction of thermodynamic phase diagrams of IDRs within a coarse-grained model defined at residue level [21]. The model includes a short-range contact potential as well as a simplified treatment of electrostatic energy. Sequence dependence is based on the short-range Miyazawa-Jernigan statistical potential [22]. The above mentioned FH approach coupled with reaction diffusion kinetics successfully models the dynamics of centrosomes but lacks specific sequence information [17]. Similarly, a multi-phase Cahn-Hilliard diffuse interface model has been used to examine RNA-protein interactions driving LLPS, demonstrating that the binding and unbinding of distinct RNA-protein complexes may lead to spatio-temporal heterogeneity within phase-separated biological condensates, in the absence of any specific sequence information [23]. Both FH and the related Overbeek and Voorn extension, that explicitly considers electrostatic effects that are required for the description of complex coacervation between oppositely charged polymers [24], are mean-field theories that best describe homo-polymers and do not consider explicit sequence-dependent effects or polymer fluctuations. The recently developed Random Phase Approximation (RPA) explicitly considers sequence patterning of charged residues, whereas fluctuations beyond the mean-field are introduced at a first order perturbation level. RPA has been successfully used to model experimental data of the DDX4 protein at a qualitative level [25]. Finally, pi-pi stacking interactions were found to involve non-aromatic as well as aromatic groups in folded globular proteins, so that a phase separation predictive algorithm was built based on pi interaction frequency [26]. The aim of this research is to set up robust and computationally efficient predictors of protein phase behavior. These predictors used as input the primary sequences of one or more proteins that may be involved in the formation of disorder bio-molecular condensates. The pathogenic mutants of proteins involved in LLPS that lead to degenerative disease has also been considered as a prediction target. In particular, the PASTA algorithm [27, 28] is one of the state-of-the-art predictors in the field of aggregation propensity into amyloid fibrils, whereas the redesigned server PASTA 2.0 [29] allows to establish the presence of cross-amyloid interactions between co-aggregating proteins. The PASTA 2.0 energy function evaluates the stability of putative cross-pairings between different sequence stretches and is based on knowledge-based statistical potentials. Along the same lines, a comprehensive knowledge-based scoring function, named BACH, was more recently developed [30]. The BACH scoring function evaluates the quality of a 3-d protein structure and allows to discriminate the proper binding modes in protein complexes [31]. It is also at the basis of a method that predicts the binding affinity of protein complexes [32].

We derived several specific knowledge-based potentials for the different kind of interactions that are believed to drive protein LLPS. The energy paramaters have been derived based on a survey of globular protein structures in the Protein Data Bank, adapting the same strategy already used for PASTA and BACH, namely the detection of residue preferences for specific interaction motifs. These included pi-pi stacking, cation-pi, and *β*-zipper formation. The latter motif was essentially the same already scored by the PASTA 2.0 energy function, but an evaluation of the biases in residue composition specific to prion-like domains [33] and low complexity protein segments [11] was required, since prion-like domains and, more generally, low complexity domains, a subset of IDRs, are found to be enriched in proteins involved in LLPS. The scoring functions built in this paper has been combined to build several predictors of the ability of a given protein or set of proteins to undergo LLPS in physiological conditions. These predictors would answer the question: “does a protein phase separate?” with a simple yes/no answer, as already proposed [26].

## Results and Discussion

In this research we construct a number of scoring functions and numerically estimate their abilities to classify a protein sequence according to its phase separating behaviour. All the protein sequences were taken from the Protein Data Bank (see supplimentary files). We used a total of 121 PhasePro (PP) proteins which are known to exhibit phase separating behaviour (*P* = 121). On the other hand we took a total of 43515 Human Proteome (HP) sequences which do not undergo phase separation (*N* = 43515). The predictability of these scoring functions are evaluated in terms of area under the curve (AUC) in receiver operator characteristics (ROC) space (see Materials and Methods), by calculating the number of true positives (*TP*), false negatives (*FN*), true negatives (*TN*) and false positives (*FP*). These scoring functions are basically different algebraic combinations of PASTA and Vernon score, where some other quantities related to the sequences were also used. We have compared our results with respect to the original Vernon score as well. The AUC denotes the ability to differentiate between the PP and HP sets. We have also evaluated the Matthews Correlation Coefficient (MCC) on the test set using the parameters optimised on the training set to study the performance of ROC analysis. In all the cases, simplex algorithm was used to maximise the AUC on the training set. The set of parameters (*α, β* or *γ*) maximising AUC are also studied.

Finally we performed 4-fold cross validation, where the ROC statistics is studied for a negative set of 43515 HP sequences by comparing them against the positive set of 121 PP sequences. The total set is divided randomly into 4 subsets. One of the subsets is taken as the training set in which the score parameters are obtained by maximising AUC and the obtained parameters are then used to evaluate AUC and MCC on the remaining subsets, known as the test set. This is repeated for all the subsets considering them as the training set and then testing the results on the corresponding test sets. We performed 100 realisations of this procedure.

In this paper we report the performances of a total of 5 scoring functions, which are defined as following:

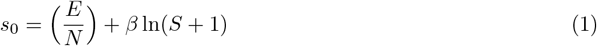

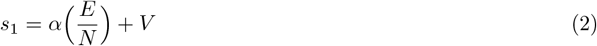

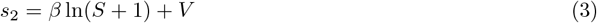

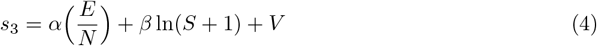

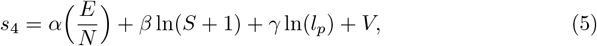

where, *E* and *V* are the PASTA and Vernon scores respectively. *N* is the number of amino acids in the sequence, *S* is the average register shift of the PASTA score over the best 5 pairings and *l*_*p*_ is the length of the best pairing. *α, β* and *γ* are the parameters which were optimised by maximising AUC on the training set and then used to find AUC and MCC on the test sets. For Vernon score we did not perform cross validation but evaluated its performance on the test set by computing the corresponding AUC and MCC for each realisation of 4-fold cross validation.

In Fig 1 we show the normalised distributions of AUC on the test set using the parameters optimised on the training set. Performance of *s*_0_ is worse than that for Vernon score. From naked-eye it clearly looks like that the distribution of AUC on test set for Vernon scores are different from all of our newly defined scores. To check statistical significance we perform one-way ANOVAs on AUC with the scoring funtion as the factor in R programming language. The results of these ANOVAs are summarised in Table 1. We found that the AUC values for all the scoring functions are statistically significant from each other. The AUC values for scores *s*_1_, *s*_2_, *s*_3_ and *s*_4_ are also statistically significant. But for the two best performing scores i.e. *s*_3_ and *s*_4_, the AUC values are not statistically significant. A series of Kolmogorov-Smirnov tests on each of the AUC distributions for different scores found that the distributions are normal for all scores (all *p*’s > 0.3) except for the score *s*_0_ (*p* = 0.003).

**Fig 1.**
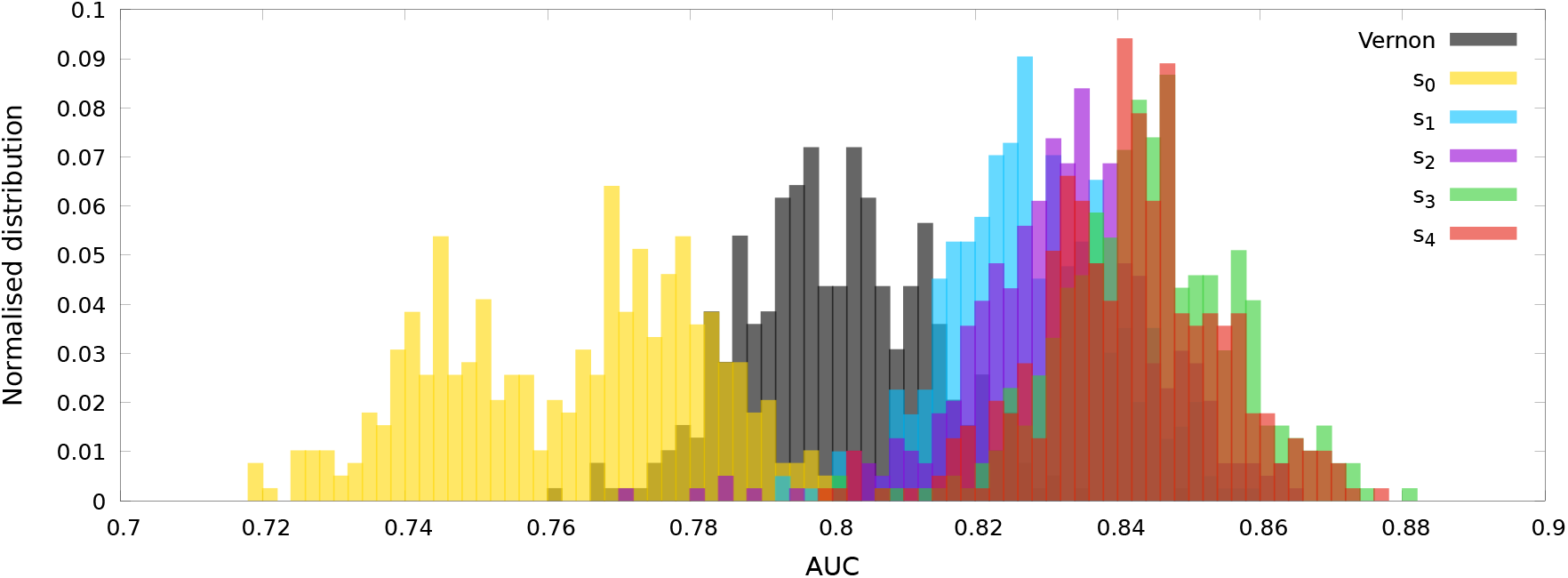
Normalised distributions for the values of AUC on the test set as obtained from the cross validation procedure for the scores *s*_0_, *s*_1_, *s*_2_, *s*_3_ and *s*_4_. The AUC values for Vernon score shown here were obtained on the test sets of cross validation. The distributions were estimated using a bin width = 0.002.

**Table 1.** Results of one-way ANOVAs to check statistical significance between AUC values on test set for several scoring functions

| Comparing<br>AUC for<br>the scores | $F$ | $p$ | $\eta^2$ |
| --- | --- | --- | --- |
| Vernon, $s_0, s_1, s_2, s_3, s_4$ | $F(5, 2349) = 1985$ | $< 10^{-6}$ | 0.808 |
| $s_1, s_2, s_3, s_4$ | $F(3, 1572) = 154.6$ | $< 10^{-6}$ | 0.228 |
| $s_3, s_4$ | $F(1, 783) = 8.453$ | 0.004 | 0.011 |

The value of AUC and MCC on the test set evaluated using the parameters optimised on the training set indicates the performance of each of the scores to segregate protein sequences according to their phase separating behaviour. In Table 2, we summarise the mean values of AUC and MCC on the test set for each of the scores, along with the optimised parameters as defined in Eqs. (1) - (5). *s*_3_ and *s*_4_ are identified as the best performing scores with AUC ≈ 0.84. Performances of the scores *s*_3_ and *s*_4_ are very similar to each other as also pointed out by the one-way ANOVA. This means that the effect of adding length of the best pairing *l*_*p*_ to *s*_3_ does not significantly change the predicting behavior.

**Table 2.**
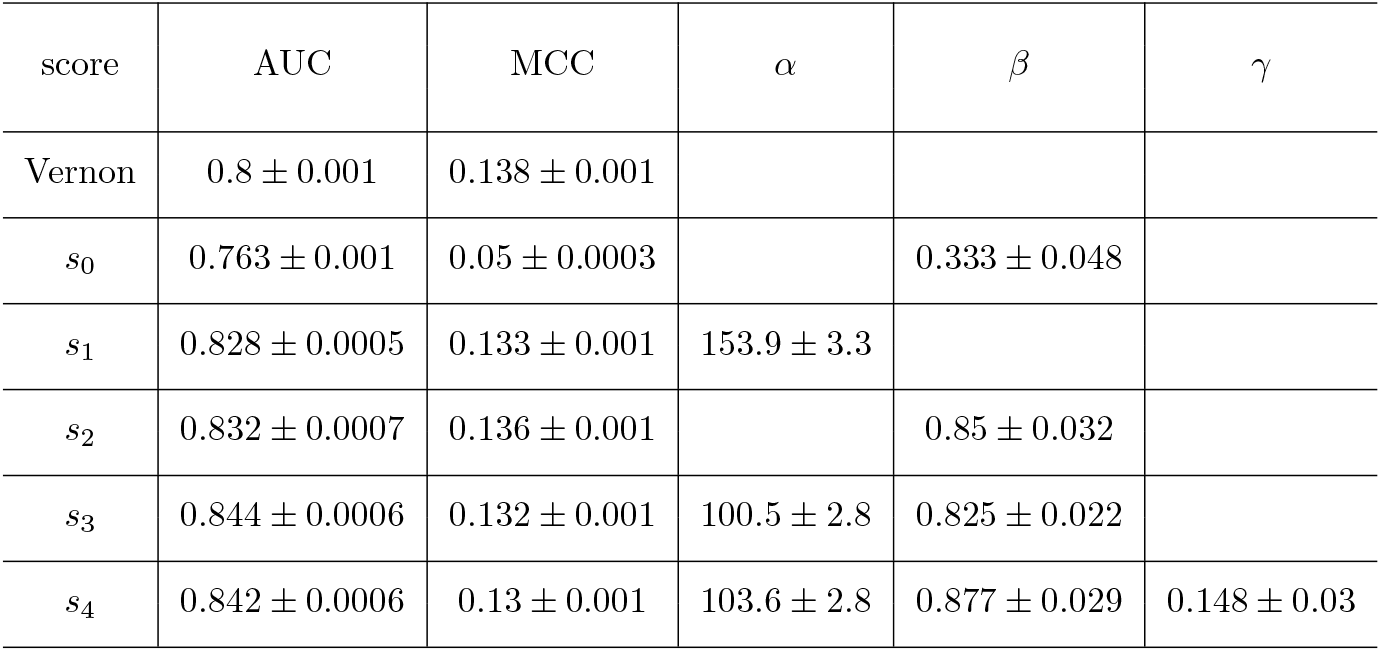
Table shows values of AUC and MCC on the test set using the parameters optimised on the training set. The optimised parameters viz. *α, β* and *γ* are also summarised here. Each of the numerical entries are mean ± standard error of the corresponding quantity.

Even after high value of AUC (∼ 0.8) in all the cases, a very low value of MCC is apparently ambiguous. However, looking carefully at the definition of MCC (see Materials and Methods), we realise that for *N* = *TN* + *FP*→ ∞ (we have *N* = 43515), with *P* = *TP* + *FN* fixed and *FP/N, TP/P* fixed, MCC actually goes to zero. Thus as a consequence of taking a larger negative set, the value of MCC on the test set remains low. We at this point realise that the study of MCC would only be meaningful when the sizes of the negative and positive sets are comparable.

We have also studied the key parameters *α, β* and *γ* in the definition of scoring functions. These parameters are optimised in the training set to produce the maximum AUC. Normalised distributions of the optimised parameters are shown in Fig 2. From Kolmogorov-Smirnov tests we found that the obtained data for *α* and *β* does not follow a normal distribution (all *p*’s < 0.01), whereas *γ* obeys a normal distribution (*p* = 0.843), as could also be seen from Fig 2.

**Fig 2.**
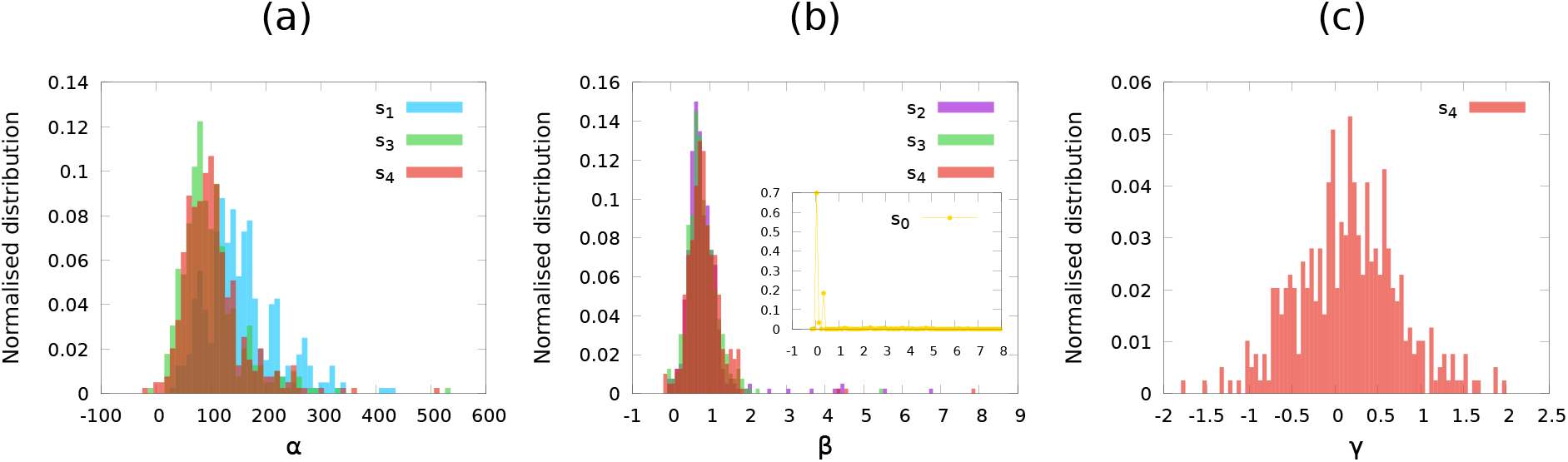
Normalised distributions of the parameters (a) *α*, (b) *β* and (c) *γ* for different scores. These parameters are optimised on the training set. *α* and *β* does not follow a normal distribution, in contrary with *γ*.

Finally, using the optimised parameters, which are summarised in Table 2, we plot the ROC curves on our entire data set for each of the predicting scores, which are shown in Fig. 3. In the vertical axis we plot the true positive rate *tpr* = *TP/*(*TP* + *FN*) and in the horizontal axis we plot the false positive rate *fpr* = *FP/*(*FP* + *TN*). Each of *s*_1_, *s*_2_, *s*_3_ and *s*_4_ acts as a better predictor than the Vernon score on our dataset. The scores *s*_3_ and *s*_4_ produces similar ROC curves with very small difference in the corresponding values of AUC as well.

**Fig 3.**
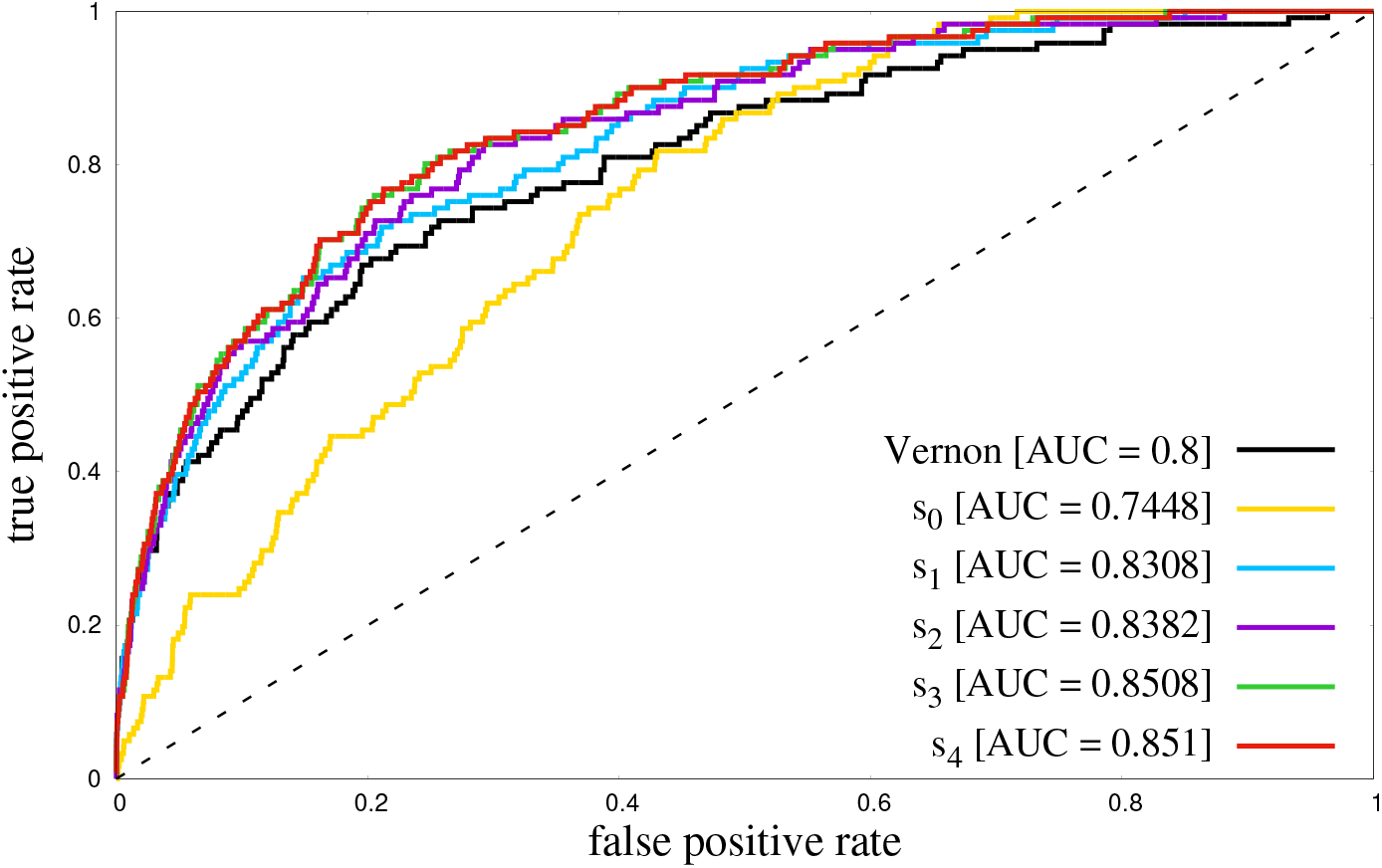
ROC curves of several predicting scores on the entire data set of 43636 protein sequences. The dashed line represents the random classifier. It is clear that *s*_1_, *s*_2_, *s*_3_ and *s*_4_ are better predicting scores than the Vernon score. This is also evident from the value of AUC.

## Materials and methods

### Register shift and PASTA score

For each of the protein sequences we use an algorithm to calculate prediction of amyloid structure aggregation (PASTA) score as has been done in [Trovato.ploscb.2006]. The estimation of PASTA score originally helped to study the role of sequence heterogeneity in driving specific aggregation into ordered self-propagating cross-*β* structures. In this approach it was assumed that only a single stretch per sequence participates in the *β*-pairing and that all other residues are not involved in aggregation and are found in a disordered noncompact conformation. Using the recipe presented in [Trovato.ploscb.2006] we calculate 5 such pairs producing the lowest PASTA scores - known as the best pairings. We also note the indices of the amino acids involved in the best pairing. For a best pair of a sequence from the index *k* to *l* registered with its parallel or anti-parallel sequence from the index *m* to *n* (*l > k* and *l* − *k* = |*m* − *n*|), the register shift *S* is defined as

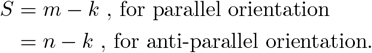

The length of the best pairing *l*_*p*_ (= *l* − *k*) was also used as a variable to define the score *s*_4_.

### Vernon score

Vernon et. al. in [vernon 2018] defined a scoring function PScore to predict liquid liquid phase separation of a protein chain using the ROC characteristics. The PScore was defined taking into account the planar pi-pi contact frequencies. The python code developed by them is available at https://doi.org/10.7554/eLife.31486.021 and we use this code to calculate the PScore which we call in our paper as the Vernon score. We originally had a total of 74811 amino acid sequences in the HP database. However, we found that the python code could not estimate a score for a total of 31296 sequences. This left us with 43515 HP chains which we use as the negative set in our machine learning analysis.

### Matthews Correlation Coefficient (MCC)

To estimate the quality of binary classification of protein sequences with respect to their phase separating behavior, we estimate the Matthews correlation coefficient (MCC). MCC is supposed to be a measure of the quality of binary classification. Numerically MCC is defined as,

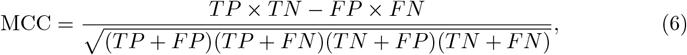

where *TP* is the number of true positives, *TN* the number of true negatives, *FP* the number of false positives and *FN* the number of false negatives. MCC basically indicates a correlation coefficient between the observed and predicted results, having a value between −1 to 1. MCC = 1 represents perfect prediction, MCC = −1 indicates a totally wrong prediction and MCC = 0 indicates a random prediction.

### *Κ* -fold Cross Validation

Cross validation is a very useful technique to asses the predictability of any predictive machine learning algorithm. This is basically a resampling method that takes into account different parts of the data as the training set and then remaining part as the test set. For a *k*-fold cross validation algorithm, the entire dataset is randomly divided into *k* parts. Choice of the size of training and test set could vary depending on the requirement. In our work, among the *k* samples, we choose *k* −1 samples as the training set and the remaining sample as the test set. During the training procedure for each of our defined scoring potentials we optimised the parameters (*α, β* or *γ*) by maximising AUC on the training set. Then using these optimised parameters we evaluate AUC (and MCC) on the test set. We repeat this process for *k* times, where in each attempt a different combination of the *k* − 1 samples as the training set. So after each step we end up with *k* set of values for the optimised parameters, maximised AUC on the training set and estimated AUC (and MCC) on the test set. Then we repeat the random division of entire dataset into *k* subsamples for a total of *N* realisations. This gives us a total of *k* × *N* set of outputs, which we then use for the statistical analysis as shown in Fig 1 and Fig 2. We used *k* = 4 and *N* = 100.

## Supporting information

A collection of all the data, code and metadata files as described in the Supplementary Materials section of the manuscript

## Supplementary materials

The following materials used for the research are publicly available at https://doi.org/10.5281/zenodo.5778299

**File 1 Phase-pro (PP) data set**. A data set of 121 protein sequences that are known to undergo phase-separation.

**File 2 Human Proteome (HP) data set**. A data set of 74811 protein sequences whose phase separation behavior is unknown or is assumed that they do not undergo phase separation.

**File 3 Pasta Code 1**. A fortran code that has been used to estimate the PASTA score for each of the protein sequences. This code detects the best 5 pairings, mentions the indices of residues involved in the pairings and denotes whether the pairing is parallel or anti-parallel.

**File 4 Pasta Code 2**. The output of the previous item is fed into this awk code to produce the following two items.

**File 5 Pasta scores for PP data set**. Output of the pasta code when applied on the PP data set, with the following details: column 1 - PASTA score, column 2 - mean PASTA score over 5 best pairings, column 3 - length of the protein sequence i.e. number of amino acid residues in the protein chain, column 4 - length of the best pair, column 5 - mean length of best 5 pairs, column 6 - register shift of the best pair, column 7 - mean register shift over best 5 pairs, column 8 - an index (*p*) indicating whether the best pair has parallel (*p* = 1) or anti-parallel (*p* = −1) orientation. The sequence of data in this file corresponds to the sequence of protein chains as in the PP data set.

**File 6 Pasta scores for HP data set**. Output of the pasta code on the HP data set, with the same details as the previous item.

**File 7 Vernon scores for the PP set**. The sequence of data in this file corresponds to the sequence of protein chains as in the PP data set.

**File 8 Vernon scores for the HP set**. Not all the 74811 HP sequences have a vernon score. Column 1 - sequence of the protein chain as in the HP data set, column 2

- vernon score of the protein chain. For the sequences which do not have a vernon score, column 2 is left blank. We had vernon scores for a total of 43515 sequences.

**File 9 Combined data file for pasta scores of 121+43515=43636 protein sequences**. The first 121 lines in this data file corresponds to the 121 protein chains in the PP data set and the next 43515 lines are for the HP sequences which do have a vernon score. This file was used as an input in the cross validation algorithm. The description of variation columns in this file is the same as File 5 and File 6.

**File 10 Combined data file for vernon scores of 121+43515=43636 protein sequences**. The arrangement of this data file is the same as the previous item. This file was also used as an input into the cross validation algorithm.

**File 11 Cross validation code**. A fortran code for *k*-fold cross validation over a number of realisations. The value *k* and the number of realisations can be changed as desired. In the training process, logistic regression was performed and the parameters of scoring functions were optimised to produce the maximum AUC. The optimised parameters were then used to calculate AUC (and MCC) on the test set.

**File 12 Output of the cross validation for score** *s*_0_. Column 1 - maximum AUC on the training set, column 2 - *β*, column 3 - AUC on the test set, column 4 - MCC on the test set.

**File 13 Output of the cross validation for score** *s*_1_. Column 1 - maximum AUC on the training set, column 2 - *α*, column 3 - AUC on the test set, column 4 - MCC on the test set.

**File 14 Output of the cross validation for score** *s*_2_. Column 1 - maximum AUC on the training set, column 2 - *β*, column 3 - AUC on the test set, column 4 - MCC on the test set.

**File 15 Output of the cross validation for score** *s*_3_. Column 1 - maximum AUC on the training set, column 2 - *α*, column 3 - *β*, column 4 - AUC on the test set, column 5 - MCC on the test set.

**File 16 Output of the cross validation for score** *s*_4_. Column 1 - maximum AUC on the training set, column 2 - *α*, column 3 - *β*, column 4 - *γ*, column 5 - AUC on the test set, column 6 - MCC on the test set.

**File 17 AUC and MCC for the vernon score on the test sets**.

## Acknowledgements

P.M. gratefully acknowledges his stay at Dipartimento di Fisica et Astronomia ‘Galileo Galilei’, University of Padova and the financial support provided (grant number: BIRD199809) to carry out this research.

