## Supplementary material for "Sequence based prediction of protein phase separation into disordered condensates using machine learning": A collection of all the data, code and metadata files as described in the Supplementary Materials section of the manuscript: file3_pasta.f

```
	program pasta
	
	implicit double precision(a-h,o-z)
	
	dimension amp(20,20),ama(20,20),is1(15000),is2(15000)
	dimension es(5),ia(5),ib(5),ic(5),id(5),ntim(3)
	character seq1*15000,seq2*15000,amin*20
	double precision,dimension(:,:),allocatable::enp,ena
	
	ent=-0.2d0

	amin='CFLWVIMHYAGPNTSRQDKE'
	
	open(70,file='pot_pasta.dat')		
	open(80,file='output')		!change
	do i=1,210
	read(70,*)n1,n2,xp,xa,xj1,xj2
c	write(*,*)n1,n2,xp,xa,xj1,xj2
	amp(n1,n2)=xp-ent
c	write(10,*)n1,n2,xp-ent,amp(n1,n2)
	amp(n2,n1)=xp-ent
	ama(n1,n2)=xa-ent
	ama(n2,n1)=xa-ent
	end do
	close(70)
	
c	do i=1,20
c	write(10,*)(amp(i,j),j=1,20)
c	end do

	open(71,file='input')		!change
	open(73,file='input1')		!change

	do ms = 1, 14811		! change for 4

	es=10000.d0	

	read(71,'(a)',end=20)seq1
20	continue
	read(73,'(a)',end=22)seq2
22	continue

	if(ms.ge.3366)then		! change


c	if(mod(ms+2,3).eq.0)then
c	write(80,*)seq1
c	end if

c	if(mod(ms+1,3).eq.0)then

	call itime(ntim)
c	write(*,"('the time is:',3i3)")ntim
	write(4444,"('the time is:',3i3)")ntim		!change

	ki=0
	do i=1,15000
	 do j=1,20
	  if(seq1(i:i)==amin(j:j)) is1(i)=j				! is1: first sequence translated into numbers
	 end do
	if(seq1(i:i).eq.'')ki=1
	if(ki.eq.1)then
	L1=i-1
	write(4444,*)'sequence',ms+60000		!change		!change
c	write(*,*)L1
	write(4444,*)L1		!change
	go to 72
	end if
	end do
72	pratik=15722
c	close(71)

c	write(80,*)(seq1(i:i),i=1,L1)
	write(80,*)'sequence',ms+60000	!change
	write(80,*)'length',L1


c	read(73,'(a)',end=30)seq2
c30 	continue

	ki=0
	do i=1,15000
	 do j=1,20
	  if(seq2(i:i)==amin(j:j)) is2(i)=j				! is2: second sequence translated into numbers
	 end do
	if(seq2(i:i).eq.'')ki=1
	if(ki.eq.1)then
	L2=i-1
	write(4444,*)L2		!change
	go to 73
	end if
	end do
73	pratik=15722
c	close(73)

	

	allocate(enp(L1,L2))
	allocate(ena(L1,L2))

	do i=1,L1	
	 do j=1,L2
	  enp(i,j)=amp(is1(i),is2(j))
	  ena(i,j)=ama(is1(i),is2(j))
	 end do
	end do
	
	
c	l=4
	do l=4,L1
c	parallel	
c	write(11,*)'parallel'
c	write(11,*)'l=',l
	 do i=1,L1
	  if((i+l-1).le.L1)then		! sliding possible
	   do j=1,L2
	    if((j+l-1).le.L2)then	! sliding possible
	    esum=0.0d0
	    i1=i
	    i2=i+l-1
	    j1=j
	    j2=j+l-1
	     do ll=0,l-1
	     esum=esum+enp(i1+ll,j1+ll)
	     end do  ! ll loop
c	     write(52,*)esum
	do ns=1,5
	 if(es(ns).ge.esum)then
	  js=6
	  do
	   js=js-1
	   if(js.eq.ns)go to 41
	    es(js)=es(js-1)
	  end do
41	  es(ns)=esum
	  ia(ns)=i1
	  ib(ns)=i2
	  ic(ns)=j1
	  id(ns)=j2
	  go to 42
	 end if
	end do
42	pratik=15722	
c	write(11,*)(seq1(k:k),k=i1,i1+l-1),'  ',(seq2(k:k),k=j1,j1+l-1),
c     &  i1,'-',i1+l-1,'&',j1,'-',j1+l-1,esum	     
c	write(11,*)ii,i1(ii),i2(ii),j1(ii),j2(ii),esum(ii)
	    end if
	   end do  ! j loop	
	  end if
	 end do	 ! i loop
	 
c	anti-parallel
c	write(11,*)'anti-parallel'
c	write(11,*)'l=',l
	do i=1,L1
	 if((i+l-1).le.L1)then		! sliding possible
	  do j=L2,1,-1
	   if((j-l+1).ge.1)then		! sliding possible
	   esum=0.0d0
	   i1=i
	   i2=i+l-1
	   j1=j
	   j2=j-l+1
	    do ll=0,l-1
	     esum=esum+ena(i1+ll,j1-ll)
	    end do  ! ll loop
c	     write(52,*)esum
	do ns=1,5
	 if(es(ns).ge.esum)then
	 js=6
	 do
	 js=js-1
	 if(js.eq.ns)go to 43
	 es(js)=es(js-1)
	 end do
43	 es(ns)=esum
	 ia(ns)=i1
	 ib(ns)=i2
	 ic(ns)=j1
	 id(ns)=j2
	 go to 44
	 end if
	end do
44	pratik=15722

c	write(11,*)(seq1(k:k),k=i1,i1+l-1),'  ',(seq2(k:k),k=j1,
c     &  j1-l+1,-1),i1,'-',i1+l-1,'&',j1,'-',j1-l+1,esum	   
c	write(11,*)ii,i1(ii),i2(ii),j1(ii),j2(ii),esum(ii)
	   end if
	  end do ! j loop
	 end if 
	end do	! i loop 
	 
	end do	! l loop
	


	do nt=1,5
	if(ic(nt).lt.id(nt))then
	write(80,11)es(nt),ia(nt),ib(nt),ic(nt),id(nt),'parallel'
11	format(f20.15,x,I6,x,I6,x,I6,x,I6,x,a)
	else
	write(80,12)es(nt),ia(nt),ib(nt),ic(nt),id(nt),'anti-parallel'
12	format(f20.15,x,I6,x,I6,x,I6,x,I6,x,a)
	end if
	end do
     
     	deallocate(ena)
     	deallocate(enp)
	end if
	end do        ! ms loop

	end program
```
