## Supplementary material for "Sequence based prediction of protein phase separation into disordered condensates using machine learning": A collection of all the data, code and metadata files as described in the Supplementary Materials section of the manuscript: file4_pasta.sh

```
awk ' BEGIN{i=0} {if(i==2){l++; eav+=$1; pairav+=$3-$2+1; p=$5-$4; if(p>0){p=1; y=$4-$2}; if(p<0){p=-1; y=$5-$2}; if(y<0.0){y=-y}; regav+=y; if(l==1){ebest=$1; pairbest=$3-$2+1; regbest=y}; if(l==5){eav/=l; pairav/=l; regav/=l; print ebest,eav,x,pairbest,pairav,regbest,regav,p; i=0}}; if($1==">"){i++;x=$5; eav=0.0; pairav=0.0; regav=0.0; l=0}} ' output > output1
```
