## Supplementary material for "Sequence based prediction of protein phase separation into disordered condensates using machine learning": A collection of all the data, code and metadata files as described in the Supplementary Materials section of the manuscript: file11_crossval.f

```
	program	cross_validation
	
	implicit double precision(a-h,o-z)
	parameter(M=43636,k=4)
	dimension npos(k),nneg(k)
	dimension na(M),nb(M)
	dimension p0(2,1),y0(2),h0(1),op0(1)
	dimension p1(2,1),y1(2),h1(1),op1(1)
	dimension p2(2,1),y2(2),h2(1),op2(1)
	dimension p3(3,2),y3(3),h3(2),op3(2)
	dimension p4(4,3),y4(4),h4(3),op4(3)		
	
	open(70,file='score0')
	open(71,file='score1')
	open(72,file='score2')
	open(73,file='score3')
	open(74,file='score4')	
	open(75,file='scorev')
	
        open(153,file='status_k4op')

        ibm=654

	do NV=1,100


	na=0
	nb=0
	npos=0
	nneg=0
	
	
	do i=1,(k-1)*M/k
c	ibm=ibm+1
100	r=ran2(ibm)
	l=r*M+1
	if(nb(l).ne.0)go to 100
	nb(l)=1
	na(i)=l
	end do
	
	iw=(k-1)*M/k
	do i=1,M
	if(nb(i).eq.0)then
	iw=iw+1
	na(iw)=i
	end if
	end do
	
	do i=0,k-1
	do j=(i*M/k)+1,(i+1)*M/k
c	if(na(j).le.121)write(11,*)na(j),i+1,'th','','positive'
c	if(na(j).gt.121)write(11,*)na(j),i+1,'th','','negative'	
	if(na(j).le.121)npos(i+1)=npos(i+1)+1
	if(na(j).gt.121)nneg(i+1)=nneg(i+1)+1
	end do
	end do
	
c	do i=1,k
c	write(*,*)npos(i),nneg(i)
c	end do

	do jk=1,k
	


	iu=((jk-1)*M/k)+1
	iv=jk*M/k
	zp=npos(jk)
	zn=nneg(jk)
	
c	write(*,*)iu,iv,zp,zn	

	ndim0=1
	np0=1
	mp0=2
	ftol0=0.00001d0	
	xmin0=10000.
	
	ndim1=1
	np1=1
	mp1=2
	ftol1=0.00001d0	
	xmin1=10000.	
	
	ndim2=1
	np2=1
	mp2=2
	ftol2=0.00001d0	
	xmin2=10000.	
	
	ndim3=2
	np3=2
	mp3=3
	ftol3=0.00001d0	
	xmin3=10000.
	
	ndim4=3
	np4=3
	mp4=4
	ftol4=0.00001d0	
	xmin4=10000.		

	do it = 1,100

	write(153,*)NV,jk,it

	r1=ran2(ibm)
	r2=ran2(ibm)
	r3=ran2(ibm)
	r4=ran2(ibm)	
	r5=ran2(ibm)
	r6=ran2(ibm)		
	r7=ran2(ibm)
	r8=ran2(ibm)
	r9=ran2(ibm)
	r10=ran2(ibm)	
	r11=ran2(ibm)
	r12=ran2(ibm)	
	r13=ran2(ibm)
	r14=ran2(ibm)
	r15=ran2(ibm)
	r16=ran2(ibm)	
	r17=ran2(ibm)
	r18=ran2(ibm)		
	r19=ran2(ibm)
	r20=ran2(ibm)
	r21=ran2(ibm)
	r22=ran2(ibm)	
	r23=ran2(ibm)
	r24=ran2(ibm)	
		

	p0(1,1)=(200.*r1) - 100.			! for score 0 
	p0(2,1)=(200.*r2) - 100.			! for score 0
	
	p1(1,1)=(200.*r3) - 100.			! for score 1
	p1(2,1)=(200.*r4) - 100.			! for score 1
	
	p2(1,1)=(200.*r5) - 100.			! for score 2
	p2(2,1)=(200.*r6) - 100.			! for score 2	
	
	p3(1,1)=(2000.*r7) - 1000.			!
	p3(1,2)=(20.*r8) - 10.				! for
	p3(2,1)=(2000.*r9) - 1000.			! score
	p3(2,2)=(20.*r10) - 10.				! 3
	p3(3,1)=(2000.*r11) - 1000.			!
	p3(3,2)=(20.*r12) - 10.				!
	
	p4(1,1)=(2000.*r13) - 1000.			!
	p4(1,2)=(20.*r14) - 10.				!
	p4(1,3)=(20.*r15) - 10.				! for
	p4(2,1)=(2000.*r16) - 1000.			!
	p4(2,2)=(20.*r17) - 10.				! score
	p4(2,3)=(20.*r18) - 10.				!
	p4(3,1)=(2000.*r19) - 1000.			! 4
	p4(3,2)=(20.*r20) - 10.				!
	p4(3,3)=(20.*r21) - 10.				!
	p4(4,1)=(2000.*r22) - 1000.			!
	p4(4,2)=(20.*r23) - 10.				!
	p4(4,3)=(20.*r24) - 10.				!
	

	do i=1,mp0
	do j=1,np0
	h0(j)=p0(i,j)
	end do
	y0(i)=area0(M,na,iu,iv,zp,zn,h0)
	end do
	
	do i=1,mp1
	do j=1,np1
	h1(j)=p1(i,j)
	end do
	y1(i)=area1(M,na,iu,iv,zp,zn,h1)
	end do	
	
	do i=1,mp2
	do j=1,np2
	h2(j)=p2(i,j)
	end do
	y2(i)=area2(M,na,iu,iv,zp,zn,h2)
	end do		
	
	do i=1,mp3
	do j=1,np3
	h3(j)=p3(i,j)
	end do
	y3(i)=area3(M,na,iu,iv,zp,zn,h3)
	end do	
	
	do i=1,mp4
	do j=1,np4
	h4(j)=p4(i,j)
	end do
	y4(i)=area4(M,na,iu,iv,zp,zn,h4)
	end do		
	

	call amoeba0(p0,y0,mp0,np0,ndim0,ftol0,M,na,iu,iv,zp,zn,
     &  area0,iter)
	call amoeba1(p1,y1,mp1,np1,ndim1,ftol1,M,na,iu,iv,zp,zn,
     &  area1,iter)
	call amoeba2(p2,y2,mp2,np2,ndim2,ftol2,M,na,iu,iv,zp,zn,
     &  area2,iter)
	call amoeba3(p3,y3,mp3,np3,ndim3,ftol3,M,na,iu,iv,zp,zn,
     &  area3,iter)     
	call amoeba4(p4,y4,mp4,np4,ndim4,ftol4,M,na,iu,iv,zp,zn,
     &  area4,iter)

c	write(*,*)'substep',jk,'iteration',it

	if(xmin0.gt.y0(1))then
	xmin0=y0(1)
	op0(1)=p0(1,1)
	end if
	
	if(xmin1.gt.y1(1))then
	xmin1=y1(1)
	op1(1)=p1(1,1)
	end if	
	
	if(xmin2.gt.y2(1))then
	xmin2=y2(1)
	op2(1)=p2(1,1)
	end if		
	
	if(xmin3.gt.y3(1))then
	xmin3=y3(1)
	op3(1)=p3(1,1)
	op3(2)=p3(1,2)
	end if			
	
	if(xmin4.gt.y4(1))then
	xmin4=y4(1)
	op4(1)=p4(1,1)
	op4(2)=p4(1,2)
	op4(3)=p4(1,3)	
	end if	
		
	end do		! it loop	
	
c	xminavg0=xminavg0+xmin0
	
	if(jk.eq.1)then
	mx=M/k + 1
	my=M
	xp=0
	xn=0
	do ixyz=2,k
	xp=xp+npos(ixyz)
	xn=xn+nneg(ixyz)
	end do
	call areamcc0(M,na,mx,my,xp,xn,op0,AC0,AM0)
	call areamcc1(M,na,mx,my,xp,xn,op1,AC1,AM1)		
	call areamcc2(M,na,mx,my,xp,xn,op2,AC2,AM2)
	call areamcc3(M,na,mx,my,xp,xn,op3,AC3,AM3)
	call areamcc4(M,na,mx,my,xp,xn,op4,AC4,AM4)
	call areamccv(M,na,mx,my,xp,xn,ACv,AMv)					
c	write(*,*)mx,my,xp,xn	
	end if
	
	if(jk.eq.k)then
	mx=1
	my=(k-1)*M/k
	xp=0
	xn=0
	do ixyz=1,k-1
	xp=xp+npos(ixyz)
	xn=xn+nneg(ixyz)
	end do
	call areamcc0(M,na,mx,my,xp,xn,op0,AC0,AM0)
	call areamcc1(M,na,mx,my,xp,xn,op1,AC1,AM1)
	call areamcc2(M,na,mx,my,xp,xn,op2,AC2,AM2)
	call areamcc3(M,na,mx,my,xp,xn,op3,AC3,AM3)					
	call areamcc4(M,na,mx,my,xp,xn,op4,AC4,AM4)
	call areamccv(M,na,mx,my,xp,xn,ACv,AMv)
c	write(*,*)mx,my,xp,xn	
	end if
	
	if((jk.gt.1).and.(jk.lt.k))then
	mx1=1
	my1=iu-1
	mx2=iv+1
	my2=M
	xp=0
	xn=0
	do ixyz=1,jk-1
	xp=xp+npos(ixyz)
	xn=xn+nneg(ixyz)	
	end do
	do ixyz=jk+1,k
	xp=xp+npos(ixyz)
	xn=xn+nneg(ixyz)	
	end do	
	call areamccc0(M,na,mx1,my1,mx2,my2,xp,xn,op0,AC0,AM0)
	call areamccc1(M,na,mx1,my1,mx2,my2,xp,xn,op1,AC1,AM1)	
	call areamccc2(M,na,mx1,my1,mx2,my2,xp,xn,op2,AC2,AM2)
	call areamccc3(M,na,mx1,my1,mx2,my2,xp,xn,op3,AC3,AM3)			
	call areamccc4(M,na,mx1,my1,mx2,my2,xp,xn,op4,AC4,AM4)
	call areamcccv(M,na,mx1,my1,mx2,my2,xp,xn,ACv,AMv)				
c	write(*,*)mx1,my1,mx2,my2,xp,xn
	end if	
	
					
	write(70,80)-xmin0,op0(1),-AC0,AM0
80	format(f20.15,x,f20.15,x,f20.15,x,f20.15)
	write(71,81)-xmin1,op1(1),-AC1,AM1
81	format(f20.15,x,f20.15,x,f20.15,x,f20.15)
	write(72,82)-xmin2,op2(1),-AC2,AM2
82	format(f20.15,x,f20.15,x,f20.15,x,f20.15)
	write(73,83)-xmin3,op3(1),op3(2),-AC3,AM3
83	format(f20.15,x,f20.15,x,f20.15,x,f20.15,x,f20.15)
	write(74,84)-xmin4,op4(1),op4(2),op4(3),-AC4,AM4
84	format(f20.15,x,f20.15,x,f20.15,x,f20.15,x,f20.15,x,f20.15)
	write(75,*)-ACv,AMv

		
	end do 		! jk loop
	
	end do 		! NV loop
	
	end program
	
	function area0(M,na,iu,iv,zp,zn,x0)				! to compute auc for the training set
	implicit double precision(a-h,o-z)				! to be fed in to simplex algo
	dimension x0(1),s0(M),indx(M),tpr(M),fpr(M)
	dimension na(M),ipro(M)

	is=0
	open(10,file='pasta')
c	open(20,file='vernon')
	do ia=1,M
	read(10,*)E,u2,aN,al,u5,u6,S,u8
c	read(20,*)ver	
	do ib=iu,iv
	if(na(ib).eq.ia)then
	is=is+1
	ipro(is)=ia
	s0(is)=x0(1)*log(S+1)+(E/aN)
	go to 200
	end if
	end do
200 	pratik=15722	
	end do
	close(10)
c	close(20)	

	ik=iv-iu+1	
	
	call indexx(ik,s0,indx)		
	
	tp=0.
	fp=0.
	
	ix=0
					
	do i=ik,1,-1
	
	ix=ix+1
	
	if(ipro(indx(i)).le.121)tp=tp+1
	if(ipro(indx(i)).gt.121)fp=fp+1
	tpr(ix)=tp/zp
	fpr(ix)=fp/zn		
	
	end do
	
	auc = tpr(1)*(fpr(2)-fpr(1)) + tpr(ik)*(fpr(ik)-fpr(ik-1))	
	do i=2,ik-1
	auc = auc + tpr(i)*(fpr(i+1)-fpr(i-1))
	end do
	area0=-auc/2.
	
	end
	
	subroutine areamcc0(M,na,mx,my,zp,zn,x0,AC0,AM0)		! to compute auc and mcc for 1st and last training sets
	implicit double precision(a-h,o-z)	
	dimension x0(1),s0(M),indx(M),tpr(M),fpr(M)
	dimension na(M),ipro(M)
	dimension xtp(M),xfp(M),xfn(M),xtn(M)

	is=0
	open(10,file='pasta')
c	open(20,file='vernon')
	do ia=1,M
	read(10,*)E,u2,aN,al,u5,u6,S,u8
c	read(20,*)ver
		
	do ib=mx,my
	if(na(ib).eq.ia)then
	is=is+1
	ipro(is)=ia
	s0(is)=x0(1)*log(S+1)+(E/aN)
	go to 200
	end if
	end do
	
	
200 	pratik=15722	
	end do
	close(10)
c	close(20)	

	ik=my-mx+1	
c	write(*,*)'for mcc',is,ik
	
	call indexx(ik,s0,indx)		
	
	tp=0.
	fp=0.

	ix=0
					
	do i=ik,1,-1
	
	ix=ix+1
	
	if(ipro(indx(i)).le.121)tp=tp+1
	if(ipro(indx(i)).gt.121)fp=fp+1
	tpr(ix)=tp/zp
	fpr(ix)=fp/zn		
	xtp(ix)=tp
	xfp(ix)=fp
	xfn(ix)=zp-tp
	xtn(ix)=zn-fp
		
	end do
	
		

	auc = tpr(1)*(fpr(2)-fpr(1)) + tpr(ik)*(fpr(ik)-fpr(ik-1))
	do i=2,ik-1
	auc = auc + tpr(i)*(fpr(i+1)-fpr(i-1))
	end do

	AC0=-auc/2.

	zmax=-10000.

	do l=1,ik

	if((xtp(l)+xfp(l)).eq.(0.0d0)) then
	xmcc = 0.
	go to 700
	end if
	
	if((xtp(l)+xfn(l)).eq.(0.0d0)) then
	xmcc = 0.
	go to 700
	end if
	
	if((xtn(l)+xfp(l)).eq.(0.0d0)) then
	xmcc = 0.
	go to 700
	end if
	
	if((xtn(l)+xfn(l)).eq.(0.0d0)) then
	xmcc = 0.
	go to 700
	end if			
	
	xmcc=((xtp(l)*xtn(l))-(xfp(l)*xfn(l)))/sqrt((xtp(l)+xfp(l))*
     &  (xtp(l)+xfn(l))*(xtn(l)+xfp(l))*(xtn(l)+xfn(l)))	
700	pratik = 15722

	if(zmax.lt.xmcc)zmax=xmcc

	end do
	
	AM0=zmax

	end	
	
	
	subroutine areamccc0(M,na,mx1,my1,mx2,my2,zp,zn,x0,AC0,AM0)			! to compute auc and mcc for intermediate training sets
	implicit double precision(a-h,o-z)	
	dimension x0(1),s0(M),indx(M),tpr(M),fpr(M)
	dimension na(M),ipro(M)
	dimension xtp(M),xfp(M),xfn(M),xtn(M)

	is=0
	open(10,file='pasta')
c	open(20,file='vernon')
	do ia=1,M
	read(10,*)E,u2,aN,al,u5,u6,S,u8
c	read(20,*)ver
		
	do ib=mx1,my1
	if(na(ib).eq.ia)then
	is=is+1
	ipro(is)=ia
	s0(is)=x0(1)*log(S+1)+(E/aN)
	go to 200
	end if
	end do
	
	do ib=mx2,my2
	if(na(ib).eq.ia)then
	is=is+1
	ipro(is)=ia
	s0(is)=x0(1)*log(S+1)+(E/aN)
	go to 200
	end if
	end do	
	
200 	pratik=15722	
	end do
	close(10)
c	close(20)	

	ik=my1-mx1+my2-mx2+2	
c	write(*,*)'comp',is,ik
	
	call indexx(ik,s0,indx)		
	
	tp=0.
	fp=0.

	ix=0
					
	do i=ik,1,-1
	
	ix=ix+1
	
	if(ipro(indx(i)).le.121)tp=tp+1
	if(ipro(indx(i)).gt.121)fp=fp+1
	tpr(ix)=tp/zp
	fpr(ix)=fp/zn		
	xtp(ix)=tp
	xfp(ix)=fp
	xfn(ix)=zp-tp
	xtn(ix)=zn-fp
		
	end do
	
		

	auc = tpr(1)*(fpr(2)-fpr(1)) + tpr(ik)*(fpr(ik)-fpr(ik-1))	
	do i=2,ik-1
	auc = auc + tpr(i)*(fpr(i+1)-fpr(i-1))
	end do

	AC0=-auc/2.

	zmax=-10000.

	do l=1,ik

	if((xtp(l)+xfp(l)).eq.(0.0d0)) then
	xmcc = 0.
	go to 700
	end if
	
	if((xtp(l)+xfn(l)).eq.(0.0d0)) then
	xmcc = 0.
	go to 700
	end if
	
	if((xtn(l)+xfp(l)).eq.(0.0d0)) then
	xmcc = 0.
	go to 700
	end if
	
	if((xtn(l)+xfn(l)).eq.(0.0d0)) then
	xmcc = 0.
	go to 700
	end if			
	
	xmcc=((xtp(l)*xtn(l))-(xfp(l)*xfn(l)))/sqrt((xtp(l)+xfp(l))*
     &  (xtp(l)+xfn(l))*(xtn(l)+xfp(l))*(xtn(l)+xfn(l)))	
700	pratik = 15722

	if(zmax.lt.xmcc)zmax=xmcc

	end do
	
	AM0=zmax

	end	
	
	function area1(M,na,iu,iv,zp,zn,x1)				! to compute auc for the training set
	implicit double precision(a-h,o-z)				! to be fed in to simplex algo
	dimension x1(1),s1(M),indx(M),tpr(M),fpr(M)
	dimension na(M),ipro(M)

	is=0
	open(10,file='pasta')
	open(20,file='vernon')
	do ia=1,M
	read(10,*)E,u2,aN,al,u5,u6,S,u8
	read(20,*)ver	
	do ib=iu,iv
	if(na(ib).eq.ia)then
	is=is+1
	ipro(is)=ia
	s1(is)=x1(1)*(E/aN) + ver
	go to 200
	end if
	end do
200 	pratik=15722	
	end do
	close(10)
	close(20)	

	ik=iv-iu+1	
	
	call indexx(ik,s1,indx)		
	
	tp=0.
	fp=0.
	
	ix=0
					
	do i=ik,1,-1
	
	ix=ix+1
	
	if(ipro(indx(i)).le.121)tp=tp+1
	if(ipro(indx(i)).gt.121)fp=fp+1
	tpr(ix)=tp/zp
	fpr(ix)=fp/zn		
	
	end do
	

	auc = tpr(1)*(fpr(2)-fpr(1)) + tpr(ik)*(fpr(ik)-fpr(ik-1))	
	do i=2,ik-1
	auc = auc + tpr(i)*(fpr(i+1)-fpr(i-1))
	end do
	area1=-auc/2.
	
	end
	
	subroutine areamcc1(M,na,mx,my,zp,zn,x1,AC1,AM1)		! to compute auc and mcc for 1st and last training sets
	implicit double precision(a-h,o-z)	
	dimension x1(1),s1(M),indx(M),tpr(M),fpr(M)
	dimension na(M),ipro(M)
	dimension xtp(M),xfp(M),xfn(M),xtn(M)

	is=0
	open(10,file='pasta')
	open(20,file='vernon')
	do ia=1,M
	read(10,*)E,u2,aN,al,u5,u6,S,u8
	read(20,*)ver
		
	do ib=mx,my
	if(na(ib).eq.ia)then
	is=is+1
	ipro(is)=ia
	s1(is)=x1(1)*(E/aN) + ver
	go to 200
	end if
	end do
	
	
200 	pratik=15722	
	end do
	close(10)
	close(20)	

	ik=my-mx+1	
c	write(*,*)'for mcc',is,ik
	
	call indexx(ik,s1,indx)		
	
	tp=0.
	fp=0.

	ix=0
					
	do i=ik,1,-1
	
	ix=ix+1
	
	if(ipro(indx(i)).le.121)tp=tp+1
	if(ipro(indx(i)).gt.121)fp=fp+1
	tpr(ix)=tp/zp
	fpr(ix)=fp/zn		
	xtp(ix)=tp
	xfp(ix)=fp
	xfn(ix)=zp-tp
	xtn(ix)=zn-fp
		
	end do
	
		

	auc = tpr(1)*(fpr(2)-fpr(1)) + tpr(ik)*(fpr(ik)-fpr(ik-1))
	do i=2,ik-1
	auc = auc + tpr(i)*(fpr(i+1)-fpr(i-1))
	end do

	AC1=-auc/2.

	zmax=-10000.

	do l=1,ik

	if((xtp(l)+xfp(l)).eq.(0.0d0)) then
	xmcc = 0.
	go to 700
	end if
	
	if((xtp(l)+xfn(l)).eq.(0.0d0)) then
	xmcc = 0.
	go to 700
	end if
	
	if((xtn(l)+xfp(l)).eq.(0.0d0)) then
	xmcc = 0.
	go to 700
	end if
	
	if((xtn(l)+xfn(l)).eq.(0.0d0)) then
	xmcc = 0.
	go to 700
	end if			
	
	xmcc=((xtp(l)*xtn(l))-(xfp(l)*xfn(l)))/sqrt((xtp(l)+xfp(l))*
     &  (xtp(l)+xfn(l))*(xtn(l)+xfp(l))*(xtn(l)+xfn(l)))	
700	pratik = 15722

	if(zmax.lt.xmcc)zmax=xmcc

	end do
	
	AM1=zmax

	end	
	
	
	subroutine areamccc1(M,na,mx1,my1,mx2,my2,zp,zn,x1,AC1,AM1)			! to compute auc and mcc for intermediate training sets
	implicit double precision(a-h,o-z)	
	dimension x1(1),s1(M),indx(M),tpr(M),fpr(M)
	dimension na(M),ipro(M)
	dimension xtp(M),xfp(M),xfn(M),xtn(M)

	is=0
	open(10,file='pasta')
	open(20,file='vernon')
	do ia=1,M
	read(10,*)E,u2,aN,al,u5,u6,S,u8
	read(20,*)ver
		
	do ib=mx1,my1
	if(na(ib).eq.ia)then
	is=is+1
	ipro(is)=ia
	s1(is)=x1(1)*(E/aN) + ver
	go to 200
	end if
	end do
	
	do ib=mx2,my2
	if(na(ib).eq.ia)then
	is=is+1
	ipro(is)=ia
	s1(is)=x1(1)*(E/aN) + ver
	go to 200
	end if
	end do	
	
200 	pratik=15722	
	end do
	close(10)
	close(20)	

	ik=my1-mx1+my2-mx2+2	
c	write(*,*)'comp',is,ik
	
	call indexx(ik,s1,indx)		
	
	tp=0.
	fp=0.

	ix=0
					
	do i=ik,1,-1
	
	ix=ix+1
	
	if(ipro(indx(i)).le.121)tp=tp+1
	if(ipro(indx(i)).gt.121)fp=fp+1
	tpr(ix)=tp/zp
	fpr(ix)=fp/zn		
	xtp(ix)=tp
	xfp(ix)=fp
	xfn(ix)=zp-tp
	xtn(ix)=zn-fp
		
	end do
	
		

	auc = tpr(1)*(fpr(2)-fpr(1)) + tpr(ik)*(fpr(ik)-fpr(ik-1))	
	do i=2,ik-1
	auc = auc + tpr(i)*(fpr(i+1)-fpr(i-1))
	end do

	AC1=-auc/2.

	zmax=-10000.

	do l=1,ik

	if((xtp(l)+xfp(l)).eq.(0.0d0)) then
	xmcc = 0.
	go to 700
	end if
	
	if((xtp(l)+xfn(l)).eq.(0.0d0)) then
	xmcc = 0.
	go to 700
	end if
	
	if((xtn(l)+xfp(l)).eq.(0.0d0)) then
	xmcc = 0.
	go to 700
	end if
	
	if((xtn(l)+xfn(l)).eq.(0.0d0)) then
	xmcc = 0.
	go to 700
	end if			
	
	xmcc=((xtp(l)*xtn(l))-(xfp(l)*xfn(l)))/sqrt((xtp(l)+xfp(l))*
     &  (xtp(l)+xfn(l))*(xtn(l)+xfp(l))*(xtn(l)+xfn(l)))	
700	pratik = 15722

	if(zmax.lt.xmcc)zmax=xmcc

	end do
	
	AM1=zmax

	end
	
	
	function area2(M,na,iu,iv,zp,zn,x2)				! to compute auc for the training set
	implicit double precision(a-h,o-z)				! to be fed in to simplex algo
	dimension x2(1),s2(M),indx(M),tpr(M),fpr(M)
	dimension na(M),ipro(M)

	is=0
	open(10,file='pasta')
	open(20,file='vernon')
	do ia=1,M
	read(10,*)E,u2,aN,al,u5,u6,S,u8
	read(20,*)ver	
	do ib=iu,iv
	if(na(ib).eq.ia)then
	is=is+1
	ipro(is)=ia
	s2(is)=x2(1)*log(S+1) + ver
	go to 200
	end if
	end do
200 	pratik=15722	
	end do
	close(10)
	close(20)	

	ik=iv-iu+1	
	
	call indexx(ik,s2,indx)		
	
	tp=0.
	fp=0.
	
	ix=0
					
	do i=ik,1,-1
	
	ix=ix+1
	
	if(ipro(indx(i)).le.121)tp=tp+1
	if(ipro(indx(i)).gt.121)fp=fp+1
	tpr(ix)=tp/zp
	fpr(ix)=fp/zn		
	
	end do
	

	auc = tpr(1)*(fpr(2)-fpr(1)) + tpr(ik)*(fpr(ik)-fpr(ik-1))	
	do i=2,ik-1
	auc = auc + tpr(i)*(fpr(i+1)-fpr(i-1))
	end do
	area2=-auc/2.
	
	end
	
	subroutine areamcc2(M,na,mx,my,zp,zn,x2,AC2,AM2)		! to compute auc and mcc for 1st and last training sets
	implicit double precision(a-h,o-z)	
	dimension x2(1),s2(M),indx(M),tpr(M),fpr(M)
	dimension na(M),ipro(M)
	dimension xtp(M),xfp(M),xfn(M),xtn(M)

	is=0
	open(10,file='pasta')
	open(20,file='vernon')
	do ia=1,M
	read(10,*)E,u2,aN,al,u5,u6,S,u8
	read(20,*)ver
		
	do ib=mx,my
	if(na(ib).eq.ia)then
	is=is+1
	ipro(is)=ia
	s2(is)=x2(1)*log(S+1) + ver
	go to 200
	end if
	end do
	
	
200 	pratik=15722	
	end do
	close(10)
	close(20)	

	ik=my-mx+1	
c	write(*,*)'for mcc',is,ik
	
	call indexx(ik,s2,indx)		
	
	tp=0.
	fp=0.

	ix=0
					
	do i=ik,1,-1
	
	ix=ix+1
	
	if(ipro(indx(i)).le.121)tp=tp+1
	if(ipro(indx(i)).gt.121)fp=fp+1
	tpr(ix)=tp/zp
	fpr(ix)=fp/zn		
	xtp(ix)=tp
	xfp(ix)=fp
	xfn(ix)=zp-tp
	xtn(ix)=zn-fp
		
	end do
	
		

	auc = tpr(1)*(fpr(2)-fpr(1)) + tpr(ik)*(fpr(ik)-fpr(ik-1))
	do i=2,ik-1
	auc = auc + tpr(i)*(fpr(i+1)-fpr(i-1))
	end do

	AC2=-auc/2.

	zmax=-10000.

	do l=1,ik

	if((xtp(l)+xfp(l)).eq.(0.0d0)) then
	xmcc = 0.
	go to 700
	end if
	
	if((xtp(l)+xfn(l)).eq.(0.0d0)) then
	xmcc = 0.
	go to 700
	end if
	
	if((xtn(l)+xfp(l)).eq.(0.0d0)) then
	xmcc = 0.
	go to 700
	end if
	
	if((xtn(l)+xfn(l)).eq.(0.0d0)) then
	xmcc = 0.
	go to 700
	end if			
	
	xmcc=((xtp(l)*xtn(l))-(xfp(l)*xfn(l)))/sqrt((xtp(l)+xfp(l))*
     &  (xtp(l)+xfn(l))*(xtn(l)+xfp(l))*(xtn(l)+xfn(l)))	
700	pratik = 15722

	if(zmax.lt.xmcc)zmax=xmcc

	end do
	
	AM2=zmax

	end	
	
	
	subroutine areamccc2(M,na,mx1,my1,mx2,my2,zp,zn,x2,AC2,AM2)			! to compute auc and mcc for intermediate training sets
	implicit double precision(a-h,o-z)	
	dimension x2(1),s2(M),indx(M),tpr(M),fpr(M)
	dimension na(M),ipro(M)
	dimension xtp(M),xfp(M),xfn(M),xtn(M)

	is=0
	open(10,file='pasta')
	open(20,file='vernon')
	do ia=1,M
	read(10,*)E,u2,aN,al,u5,u6,S,u8
	read(20,*)ver
		
	do ib=mx1,my1
	if(na(ib).eq.ia)then
	is=is+1
	ipro(is)=ia
	s2(is)=x2(1)*log(S+1) + ver
	go to 200
	end if
	end do
	
	do ib=mx2,my2
	if(na(ib).eq.ia)then
	is=is+1
	ipro(is)=ia
	s2(is)=x2(1)*log(S+1) + ver
	go to 200
	end if
	end do	
	
200 	pratik=15722	
	end do
	close(10)
	close(20)	

	ik=my1-mx1+my2-mx2+2	
c	write(*,*)'comp',is,ik
	
	call indexx(ik,s2,indx)		
	
	tp=0.
	fp=0.

	ix=0
					
	do i=ik,1,-1
	
	ix=ix+1
	
	if(ipro(indx(i)).le.121)tp=tp+1
	if(ipro(indx(i)).gt.121)fp=fp+1
	tpr(ix)=tp/zp
	fpr(ix)=fp/zn		
	xtp(ix)=tp
	xfp(ix)=fp
	xfn(ix)=zp-tp
	xtn(ix)=zn-fp
		
	end do
	
		

	auc = tpr(1)*(fpr(2)-fpr(1)) + tpr(ik)*(fpr(ik)-fpr(ik-1))	
	do i=2,ik-1
	auc = auc + tpr(i)*(fpr(i+1)-fpr(i-1))
	end do

	AC2=-auc/2.

	zmax=-10000.

	do l=1,ik

	if((xtp(l)+xfp(l)).eq.(0.0d0)) then
	xmcc = 0.
	go to 700
	end if
	
	if((xtp(l)+xfn(l)).eq.(0.0d0)) then
	xmcc = 0.
	go to 700
	end if
	
	if((xtn(l)+xfp(l)).eq.(0.0d0)) then
	xmcc = 0.
	go to 700
	end if
	
	if((xtn(l)+xfn(l)).eq.(0.0d0)) then
	xmcc = 0.
	go to 700
	end if			
	
	xmcc=((xtp(l)*xtn(l))-(xfp(l)*xfn(l)))/sqrt((xtp(l)+xfp(l))*
     &  (xtp(l)+xfn(l))*(xtn(l)+xfp(l))*(xtn(l)+xfn(l)))	
700	pratik = 15722

	if(zmax.lt.xmcc)zmax=xmcc

	end do
	
	AM2=zmax

	end		
	
	
	function area3(M,na,iu,iv,zp,zn,x3)				! to compute auc for the training set
	implicit double precision(a-h,o-z)				! to be fed in to simplex algo
	dimension x3(2),s3(M),indx(M),tpr(M),fpr(M)
	dimension na(M),ipro(M)
	

	is=0
	open(10,file='pasta')
	open(20,file='vernon')
	do ia=1,M
	read(10,*)E,u2,aN,al,u5,u6,S,u8
	read(20,*)ver	
	do ib=iu,iv
	if(na(ib).eq.ia)then
	is=is+1
	ipro(is)=ia
	s3(is)= x3(2)*log(S+1) + x3(1)*(E/aN) + ver
	go to 200
	end if
	end do
200 	pratik=15722	
	end do
	close(10)
	close(20)	

	ik=iv-iu+1	
	
	call indexx(ik,s3,indx)		
	
	tp=0.
	fp=0.
	
	ix=0
					
	do i=ik,1,-1
	
	ix=ix+1
	
	if(ipro(indx(i)).le.121)tp=tp+1
	if(ipro(indx(i)).gt.121)fp=fp+1
	tpr(ix)=tp/zp
	fpr(ix)=fp/zn		
	
	end do
	

	auc = tpr(1)*(fpr(2)-fpr(1)) + tpr(ik)*(fpr(ik)-fpr(ik-1))	
	do i=2,ik-1
	auc = auc + tpr(i)*(fpr(i+1)-fpr(i-1))
	end do
	area3=-auc/2.
	
	end
	
	subroutine areamcc3(M,na,mx,my,zp,zn,x3,AC3,AM3)		! to compute auc and mcc for 1st and last training sets
	implicit double precision(a-h,o-z)	
	dimension x3(2),s3(M),indx(M),tpr(M),fpr(M)
	dimension na(M),ipro(M)
	dimension xtp(M),xfp(M),xfn(M),xtn(M)

	is=0
	open(10,file='pasta')
	open(20,file='vernon')
	do ia=1,M
	read(10,*)E,u2,aN,al,u5,u6,S,u8
	read(20,*)ver
		
	do ib=mx,my
	if(na(ib).eq.ia)then
	is=is+1
	ipro(is)=ia
	s3(is)= x3(2)*log(S+1) + x3(1)*(E/aN) + ver
	go to 200
	end if
	end do
	
	
200 	pratik=15722	
	end do
	close(10)
	close(20)	

	ik=my-mx+1	
c	write(*,*)'for mcc',is,ik
	
	call indexx(ik,s3,indx)		
	
	tp=0.
	fp=0.

	ix=0
					
	do i=ik,1,-1
	
	ix=ix+1
	
	if(ipro(indx(i)).le.121)tp=tp+1
	if(ipro(indx(i)).gt.121)fp=fp+1
	tpr(ix)=tp/zp
	fpr(ix)=fp/zn		
	xtp(ix)=tp
	xfp(ix)=fp
	xfn(ix)=zp-tp
	xtn(ix)=zn-fp
	
c	write(777,*)xtp(ix),xfp(ix),xfn(ix),xtn(ix)
		
	end do
	
		

	auc = tpr(1)*(fpr(2)-fpr(1)) + tpr(ik)*(fpr(ik)-fpr(ik-1))
	do i=2,ik-1
	auc = auc + tpr(i)*(fpr(i+1)-fpr(i-1))
	end do

	AC3=-auc/2.

	zmax=-10000.

	do l=1,ik

	if((xtp(l)+xfp(l)).eq.(0.0d0)) then
	xmcc = 0.
	go to 700
	end if
	
	if((xtp(l)+xfn(l)).eq.(0.0d0)) then
	xmcc = 0.
	go to 700
	end if
	
	if((xtn(l)+xfp(l)).eq.(0.0d0)) then
	xmcc = 0.
	go to 700
	end if
	
	if((xtn(l)+xfn(l)).eq.(0.0d0)) then
	xmcc = 0.
	go to 700
	end if			
	
	xmcc=((xtp(l)*xtn(l))-(xfp(l)*xfn(l)))/sqrt((xtp(l)+xfp(l))*
     &  (xtp(l)+xfn(l))*(xtn(l)+xfp(l))*(xtn(l)+xfn(l)))	
700	pratik = 15722

	if(zmax.lt.xmcc)zmax=xmcc

	end do
	
	AM3=zmax

	end	
	
	
	subroutine areamccc3(M,na,mx1,my1,mx2,my2,zp,zn,x3,AC3,AM3)			! to compute auc and mcc for intermediate training sets
	implicit double precision(a-h,o-z)	
	dimension x3(2),s3(M),indx(M),tpr(M),fpr(M)
	dimension na(M),ipro(M)
	dimension xtp(M),xfp(M),xfn(M),xtn(M)

	is=0
	open(10,file='pasta')
	open(20,file='vernon')
	do ia=1,M
	read(10,*)E,u2,aN,al,u5,u6,S,u8
	read(20,*)ver
		
	do ib=mx1,my1
	if(na(ib).eq.ia)then
	is=is+1
	ipro(is)=ia
	s3(is)= x3(2)*log(S+1) + x3(1)*(E/aN) + ver
	go to 200
	end if
	end do
	
	do ib=mx2,my2
	if(na(ib).eq.ia)then
	is=is+1
	ipro(is)=ia
	s3(is)= x3(2)*log(S+1) + x3(1)*(E/aN) + ver
	go to 200
	end if
	end do	
	
200 	pratik=15722	
	end do
	close(10)
	close(20)	

	ik=my1-mx1+my2-mx2+2	
c	write(*,*)'comp',is,ik
	
	call indexx(ik,s3,indx)		
	
	tp=0.
	fp=0.

	ix=0
					
	do i=ik,1,-1
	
	ix=ix+1
	
	if(ipro(indx(i)).le.121)tp=tp+1
	if(ipro(indx(i)).gt.121)fp=fp+1
	tpr(ix)=tp/zp
	fpr(ix)=fp/zn		
	xtp(ix)=tp
	xfp(ix)=fp
	xfn(ix)=zp-tp
	xtn(ix)=zn-fp
		
	end do
	
		

	auc = tpr(1)*(fpr(2)-fpr(1)) + tpr(ik)*(fpr(ik)-fpr(ik-1))	
	do i=2,ik-1
	auc = auc + tpr(i)*(fpr(i+1)-fpr(i-1))
	end do

	AC3=-auc/2.

	zmax=-10000.

	do l=1,ik

	if((xtp(l)+xfp(l)).eq.(0.0d0)) then
	xmcc = 0.
	go to 700
	end if
	
	if((xtp(l)+xfn(l)).eq.(0.0d0)) then
	xmcc = 0.
	go to 700
	end if
	
	if((xtn(l)+xfp(l)).eq.(0.0d0)) then
	xmcc = 0.
	go to 700
	end if
	
	if((xtn(l)+xfn(l)).eq.(0.0d0)) then
	xmcc = 0.
	go to 700
	end if			
	
	xmcc=((xtp(l)*xtn(l))-(xfp(l)*xfn(l)))/sqrt((xtp(l)+xfp(l))*
     &  (xtp(l)+xfn(l))*(xtn(l)+xfp(l))*(xtn(l)+xfn(l)))	
700	pratik = 15722

	if(zmax.lt.xmcc)zmax=xmcc

	end do
	
	AM3=zmax

	end	
	
	
	function area4(M,na,iu,iv,zp,zn,x4)				! to compute auc for the training set
	implicit double precision(a-h,o-z)				! to be fed in to simplex algo
	dimension x4(3),s4(M),indx(M),tpr(M),fpr(M)
	dimension na(M),ipro(M)

	is=0
	open(10,file='pasta')
	open(20,file='vernon')
	do ia=1,M
	read(10,*)E,u2,aN,al,u5,u6,S,u8
	read(20,*)ver	
	do ib=iu,iv
	if(na(ib).eq.ia)then
	is=is+1
	ipro(is)=ia
	s4(is)=x4(2)*log(S+1) + x4(1)*(E/aN) + x4(3)*log(al) + ver
	go to 200
	end if
	end do
200 	pratik=15722	
	end do
	close(10)
	close(20)	

	ik=iv-iu+1	
	
	call indexx(ik,s4,indx)		
	
	tp=0.
	fp=0.
	
	ix=0
					
	do i=ik,1,-1
	
	ix=ix+1
	
	if(ipro(indx(i)).le.121)tp=tp+1
	if(ipro(indx(i)).gt.121)fp=fp+1
	tpr(ix)=tp/zp
	fpr(ix)=fp/zn		
	
	end do
	

	auc = tpr(1)*(fpr(2)-fpr(1)) + tpr(ik)*(fpr(ik)-fpr(ik-1))	
	do i=2,ik-1
	auc = auc + tpr(i)*(fpr(i+1)-fpr(i-1))
	end do
	area4=-auc/2.
	
	end
	
	subroutine areamcc4(M,na,mx,my,zp,zn,x4,AC4,AM4)		! to compute auc and mcc for 1st and last training sets
	implicit double precision(a-h,o-z)	
	dimension x4(3),s4(M),indx(M),tpr(M),fpr(M)
	dimension na(M),ipro(M)
	dimension xtp(M),xfp(M),xfn(M),xtn(M)

	is=0
	open(10,file='pasta')
	open(20,file='vernon')
	do ia=1,M
	read(10,*)E,u2,aN,al,u5,u6,S,u8
	read(20,*)ver
		
	do ib=mx,my
	if(na(ib).eq.ia)then
	is=is+1
	ipro(is)=ia
	s4(is)=x4(2)*log(S+1) + x4(1)*(E/aN) + x4(3)*log(al) + ver
	go to 200
	end if
	end do
	
	
200 	pratik=15722	
	end do
	close(10)
	close(20)	

	ik=my-mx+1	
c	write(*,*)'for mcc',is,ik
	
	call indexx(ik,s4,indx)		
	
	tp=0.
	fp=0.

	ix=0
					
	do i=ik,1,-1
	
	ix=ix+1
	
	if(ipro(indx(i)).le.121)tp=tp+1
	if(ipro(indx(i)).gt.121)fp=fp+1
	tpr(ix)=tp/zp
	fpr(ix)=fp/zn		
	xtp(ix)=tp
	xfp(ix)=fp
	xfn(ix)=zp-tp
	xtn(ix)=zn-fp
		
	end do
	
		

	auc = tpr(1)*(fpr(2)-fpr(1)) + tpr(ik)*(fpr(ik)-fpr(ik-1))
	do i=2,ik-1
	auc = auc + tpr(i)*(fpr(i+1)-fpr(i-1))
	end do

	AC4=-auc/2.

	zmax=-10000.

	do l=1,ik

	if((xtp(l)+xfp(l)).eq.(0.0d0)) then
	xmcc = 0.
	go to 700
	end if
	
	if((xtp(l)+xfn(l)).eq.(0.0d0)) then
	xmcc = 0.
	go to 700
	end if
	
	if((xtn(l)+xfp(l)).eq.(0.0d0)) then
	xmcc = 0.
	go to 700
	end if
	
	if((xtn(l)+xfn(l)).eq.(0.0d0)) then
	xmcc = 0.
	go to 700
	end if			
	
	xmcc=((xtp(l)*xtn(l))-(xfp(l)*xfn(l)))/sqrt((xtp(l)+xfp(l))*
     &  (xtp(l)+xfn(l))*(xtn(l)+xfp(l))*(xtn(l)+xfn(l)))	
700	pratik = 15722

	if(zmax.lt.xmcc)zmax=xmcc

	end do
	
	AM4=zmax

	end	
	
	
	subroutine areamccc4(M,na,mx1,my1,mx2,my2,zp,zn,x4,AC4,AM4)			! to compute auc and mcc for intermediate training sets
	implicit double precision(a-h,o-z)	
	dimension x4(3),s4(M),indx(M),tpr(M),fpr(M)
	dimension na(M),ipro(M)
	dimension xtp(M),xfp(M),xfn(M),xtn(M)

	is=0
	open(10,file='pasta')
	open(20,file='vernon')
	do ia=1,M
	read(10,*)E,u2,aN,al,u5,u6,S,u8
	read(20,*)ver
		
	do ib=mx1,my1
	if(na(ib).eq.ia)then
	is=is+1
	ipro(is)=ia
	s4(is)=x4(2)*log(S+1) + x4(1)*(E/aN) + x4(3)*log(al) + ver
	go to 200
	end if
	end do
	
	do ib=mx2,my2
	if(na(ib).eq.ia)then
	is=is+1
	ipro(is)=ia
	s4(is)=x4(2)*log(S+1) + x4(1)*(E/aN) + x4(3)*log(al) + ver
	go to 200
	end if
	end do	
	
200 	pratik=15722	
	end do
	close(10)
	close(20)	

	ik=my1-mx1+my2-mx2+2	
c	write(*,*)'comp',is,ik
	
	call indexx(ik,s4,indx)		
	
	tp=0.
	fp=0.

	ix=0
					
	do i=ik,1,-1
	
	ix=ix+1
	
	if(ipro(indx(i)).le.121)tp=tp+1
	if(ipro(indx(i)).gt.121)fp=fp+1
	tpr(ix)=tp/zp
	fpr(ix)=fp/zn		
	xtp(ix)=tp
	xfp(ix)=fp
	xfn(ix)=zp-tp
	xtn(ix)=zn-fp
		
	end do
	
		

	auc = tpr(1)*(fpr(2)-fpr(1)) + tpr(ik)*(fpr(ik)-fpr(ik-1))	
	do i=2,ik-1
	auc = auc + tpr(i)*(fpr(i+1)-fpr(i-1))
	end do

	AC4=-auc/2.

	zmax=-10000.

	do l=1,ik

	if((xtp(l)+xfp(l)).eq.(0.0d0)) then
	xmcc = 0.
	go to 700
	end if
	
	if((xtp(l)+xfn(l)).eq.(0.0d0)) then
	xmcc = 0.
	go to 700
	end if
	
	if((xtn(l)+xfp(l)).eq.(0.0d0)) then
	xmcc = 0.
	go to 700
	end if
	
	if((xtn(l)+xfn(l)).eq.(0.0d0)) then
	xmcc = 0.
	go to 700
	end if			
	
	xmcc=((xtp(l)*xtn(l))-(xfp(l)*xfn(l)))/sqrt((xtp(l)+xfp(l))*
     &  (xtp(l)+xfn(l))*(xtn(l)+xfp(l))*(xtn(l)+xfn(l)))	
700	pratik = 15722

	if(zmax.lt.xmcc)zmax=xmcc

	end do
	
	AM4=zmax

	end		

	
	subroutine areamccv(M,na,mx,my,zp,zn,ACv,AMv)		! to compute auc and mcc for 1st and last training sets
	implicit double precision(a-h,o-z)	
	dimension v(M),indx(M),tpr(M),fpr(M)
	dimension na(M),ipro(M)
	dimension xtp(M),xfp(M),xfn(M),xtn(M)

	is=0
c	open(10,file='pasta')
	open(20,file='vernon')
	do ia=1,M
c	read(10,*)E,u2,aN,al,u5,u6,S,u8
	read(20,*)ver
		
	do ib=mx,my
	if(na(ib).eq.ia)then
	is=is+1
	ipro(is)=ia
	v(is)=ver
	go to 200
	end if
	end do
	
	
200 	pratik=15722	
	end do
c	close(10)
	close(20)	

	ik=my-mx+1	
c	write(*,*)'for mcc',is,ik
	
	call indexx(ik,v,indx)		
	
	tp=0.
	fp=0.

	ix=0
					
	do i=ik,1,-1
	
	ix=ix+1
	
	if(ipro(indx(i)).le.121)tp=tp+1
	if(ipro(indx(i)).gt.121)fp=fp+1
	tpr(ix)=tp/zp
	fpr(ix)=fp/zn		
	xtp(ix)=tp
	xfp(ix)=fp
	xfn(ix)=zp-tp
	xtn(ix)=zn-fp

c	write(777,153)xtp(ix),xfp(ix),xfn(ix),xtn(ix)
c153	format(f20.15,x,f20.15,x,f20.15,x,f20.15)
		
	end do
	
		

	auc = tpr(1)*(fpr(2)-fpr(1)) + tpr(ik)*(fpr(ik)-fpr(ik-1))
	do i=2,ik-1
	auc = auc + tpr(i)*(fpr(i+1)-fpr(i-1))
	end do

	ACv=-auc/2.

	zmax=-10000.

	do l=1,ik

	if((xtp(l)+xfp(l)).eq.(0.0d0)) then
	xmcc = 0.
	go to 700
	end if
	
	if((xtp(l)+xfn(l)).eq.(0.0d0)) then
	xmcc = 0.
	go to 700
	end if
	
	if((xtn(l)+xfp(l)).eq.(0.0d0)) then
	xmcc = 0.
	go to 700
	end if
	
	if((xtn(l)+xfn(l)).eq.(0.0d0)) then
	xmcc = 0.
	go to 700
	end if			
	
	xmcc=((xtp(l)*xtn(l))-(xfp(l)*xfn(l)))/sqrt((xtp(l)+xfp(l))*
     &  (xtp(l)+xfn(l))*(xtn(l)+xfp(l))*(xtn(l)+xfn(l)))	
700	pratik = 15722

	if(zmax.lt.xmcc)zmax=xmcc

	end do
	
	AMv=zmax

	end	
	
	
	subroutine areamcccv(M,na,mx1,my1,mx2,my2,zp,zn,ACv,AMv)			! to compute auc and mcc for intermediate training sets
	implicit double precision(a-h,o-z)	
	dimension v(M),indx(M),tpr(M),fpr(M)
	dimension na(M),ipro(M)
	dimension xtp(M),xfp(M),xfn(M),xtn(M)

	is=0
c	open(10,file='pasta')
	open(20,file='vernon')
	do ia=1,M
c	read(10,*)E,u2,aN,al,u5,u6,S,u8
	read(20,*)ver
		
	do ib=mx1,my1
	if(na(ib).eq.ia)then
	is=is+1
	ipro(is)=ia
	v(is)=ver
	go to 200
	end if
	end do
	
	do ib=mx2,my2
	if(na(ib).eq.ia)then
	is=is+1
	ipro(is)=ia
	v(is)=ver
	go to 200
	end if
	end do	
	
200 	pratik=15722	
	end do
c	close(10)
	close(20)	

	ik=my1-mx1+my2-mx2+2	
c	write(*,*)'comp',is,ik
	
	call indexx(ik,v,indx)		
	
	tp=0.
	fp=0.

	ix=0
					
	do i=ik,1,-1
	
	ix=ix+1
	
	if(ipro(indx(i)).le.121)tp=tp+1
	if(ipro(indx(i)).gt.121)fp=fp+1
	tpr(ix)=tp/zp
	fpr(ix)=fp/zn		
	xtp(ix)=tp
	xfp(ix)=fp
	xfn(ix)=zp-tp
	xtn(ix)=zn-fp
		
	end do
	
		

	auc = tpr(1)*(fpr(2)-fpr(1)) + tpr(ik)*(fpr(ik)-fpr(ik-1))	
	do i=2,ik-1
	auc = auc + tpr(i)*(fpr(i+1)-fpr(i-1))
	end do

	ACv=-auc/2.

	zmax=-10000.

	do l=1,ik

	if((xtp(l)+xfp(l)).eq.(0.0d0)) then
	xmcc = 0.
	go to 700
	end if
	
	if((xtp(l)+xfn(l)).eq.(0.0d0)) then
	xmcc = 0.
	go to 700
	end if
	
	if((xtn(l)+xfp(l)).eq.(0.0d0)) then
	xmcc = 0.
	go to 700
	end if
	
	if((xtn(l)+xfn(l)).eq.(0.0d0)) then
	xmcc = 0.
	go to 700
	end if			
	
	xmcc=((xtp(l)*xtn(l))-(xfp(l)*xfn(l)))/sqrt((xtp(l)+xfp(l))*
     &  (xtp(l)+xfn(l))*(xtn(l)+xfp(l))*(xtn(l)+xfn(l)))	
700	pratik = 15722

	if(zmax.lt.xmcc)zmax=xmcc

	end do
	
	AMv=zmax

	end			
	
	
	SUBROUTINE indexx(n,arr,indx)
	implicit double precision(a-h,o-z)

	dimension arr(n),indx(n)
	PARAMETER (M=7,NSTACK=50)	
	INTEGER istack(NSTACK)
c	REAL a
	
c	write(*,*)'hello index'	
	
	do j=1,n
	indx(j)=j
	enddo 
	jstack=0
	l=1
	ir=n
1	if(ir-l.lt.M)then
	do j=l+1,ir
	indxt=indx(j)
	a=arr(indxt)
	do i=j-1,l,-1
	if(arr(indx(i)).le.a)goto 2
	indx(i+1)=indx(i)
	enddo
	i=l-1
2	indx(i+1)=indxt
	enddo
	if(jstack.eq.0)return
	ir=istack(jstack)
	l=istack(jstack-1)
	jstack=jstack-2
	else
	k=(l+ir)/2
	itemp=indx(k)
	indx(k)=indx(l+1)
	indx(l+1)=itemp
	if(arr(indx(l)).gt.arr(indx(ir)))then
	itemp=indx(l)
	indx(l)=indx(ir)
	indx(ir)=itemp
	endif
	if(arr(indx(l+1)).gt.arr(indx(ir)))then
	itemp=indx(l+1)
	indx(l+1)=indx(ir)
	indx(ir)=itemp
	endif
	if(arr(indx(l)).gt.arr(indx(l+1)))then
	itemp=indx(l)
	indx(l)=indx(l+1)
	indx(l+1)=itemp
	endif
	i=l+1
	j=ir
	indxt=indx(l+1)
	a=arr(indxt)
3 	continue
	i=i+1
	if(arr(indx(i)).lt.a)goto 3
4	continue
	j=j-1
	if(arr(indx(j)).gt.a)goto 4
	if(j.lt.i)goto 5
	itemp=indx(i)
	indx(i)=indx(j)
	indx(j)=itemp
	goto 3
5	indx(l+1)=indx(j)
	indx(j)=indxt
	jstack=jstack+2
	if(jstack.gt.NSTACK)then
	write(*,*) 'NSTACK too small in indexx'
	stop
	end if
	if(ir-i+1.ge.j-l)then
	istack(jstack)=ir
	istack(jstack-1)=i
	ir=j-1
	else
	istack(jstack)=j-1
	istack(jstack-1)=l
	l=i
	endif
	endif
	goto 1
	END
	
	SUBROUTINE amoeba0(p,y,mp,np,ndim,ftol,M,na,iu,iv,zp,zn,
     &  area0,iter)
	implicit double precision(a-h,o-z)
	dimension p(mp,np),y(mp),na(M)
	PARAMETER (NMAX=20,ITMAX=5000,TINY=1.e-10) 
	EXTERNAL area0	
	
	double precision psum(NMAX)
	
	iter=0
1 	do n=1,ndim
	 sum=0.
	 do m1=1,ndim+1
	  sum=sum+p(m1,n)
	 end do
	 psum(n)=sum
	end do

2 	ilo=1
	if (y(1).gt.y(2)) then
	 ihi=1
	 inhi=2
	else
	 ihi=2
	 inhi=1
	end if

	do i=1,ndim+1
	 if(y(i).le.y(ilo)) ilo=i
	 if(y(i).gt.y(ihi)) then
	  inhi=ihi
	  ihi=i
	 else if(y(i).gt.y(inhi)) then
	 if(i.ne.ihi) inhi=i
	 end if
	end do
	

	rtol=2.*abs(y(ihi)-y(ilo))/(abs(y(ihi))+abs(y(ilo))+TINY)
c	write(*,*)rtol

	if (rtol.lt.ftol) then 
	 swap=y(1)
	 y(1)=y(ilo)
	 y(ilo)=swap
	 do n=1,ndim
	  swap=p(1,n)
	  p(1,n)=p(ilo,n)
	  p(ilo,n)=swap
	 end do
	 return
	end if

	if (iter.ge.ITMAX)then
	write(*,*)'exceeds'
	stop !’ITMAX exceeded in amoeba’
	end if

	iter=iter+2

	ytry=amotry0(p,y,psum,mp,np,ndim,M,na,iu,iv,zp,zn,	
     &  area0,ihi,-1.0d0)	
	if (ytry.le.y(ilo)) then
	 ytry=amotry0(p,y,psum,mp,np,ndim,M,na,iu,iv,zp,zn,	
     &   area0,ihi,2.0d0)	
	else if (ytry.ge.y(inhi)) then
	 ysave=y(ihi)
	 ytry=amotry0(p,y,psum,mp,np,ndim,M,na,iu,iv,zp,zn,	
     &   area0,ihi,0.5d0)	
	 if (ytry.ge.ysave) then
	  do i=1,ndim+1
	   if(i.ne.ilo)then
	    do j=1,ndim
	     psum(j)=0.5d0*(p(i,j)+p(ilo,j))
	     p(i,j)=psum(j)
	    end do
	   y(i)=area0(M,na,iu,iv,zp,zn,psum)	
	   end if
	  end do
	  iter=iter+ndim
	  go to 1
	 end if
	else
	 iter=iter-1
	end if
	go to 2
	END

	FUNCTION amotry0(p,y,psum,mp,np,ndim,M,na,iu,iv,zp,zn,
     &  area0,ihi,fac)
	implicit double precision(a-h,o-z)
	dimension p(mp,np),psum(np),y(mp),na(M)
	PARAMETER (NMAX=20)
	EXTERNAL area0
	dimension ptry(NMAX)

	fac1=(1.0d0-fac)/ndim
	fac2=fac1-fac
	do j=1,ndim
	ptry(j)=psum(j)*fac1-p(ihi,j)*fac2
	end do
	ytry=area0(M,na,iu,iv,zp,zn,ptry) 	
	if (ytry.lt.y(ihi)) then
	y(ihi)=ytry
	do j=1,ndim
	psum(j)=psum(j)-p(ihi,j)+ptry(j)
	p(ihi,j)=ptry(j)
	end do
	end if
	amotry0=ytry	
	return
	END

	SUBROUTINE amoeba1(p,y,mp,np,ndim,ftol,M,na,iu,iv,zp,zn,	
     &  area1,iter)	
	implicit double precision(a-h,o-z)
	dimension p(mp,np),y(mp),na(M)
	PARAMETER (NMAX=20,ITMAX=5000,TINY=1.e-10) 
	EXTERNAL area1	

	double precision psum(NMAX)	

	iter=0
1 	do n=1,ndim
	 sum=0. 
	 do m1=1,ndim+1
	  sum=sum+p(m1,n)
	 end do
	 psum(n)=sum
	end do

2 	ilo=1
	if (y(1).gt.y(2)) then 
	 ihi=1
	 inhi=2
	else
	 ihi=2
	 inhi=1
	end if

	do i=1,ndim+1
	 if(y(i).le.y(ilo)) ilo=i
	 if(y(i).gt.y(ihi)) then
	  inhi=ihi
	  ihi=i
	 else if(y(i).gt.y(inhi)) then
	 if(i.ne.ihi) inhi=i
	 end if
	end do


	rtol=2.*abs(y(ihi)-y(ilo))/(abs(y(ihi))+abs(y(ilo))+TINY)
c	write(*,*)rtol

	if (rtol.lt.ftol) then 
	 swap=y(1)
	 y(1)=y(ilo)
	 y(ilo)=swap
	 do n=1,ndim
	  swap=p(1,n)
	  p(1,n)=p(ilo,n)
	  p(ilo,n)=swap
	 end do
	 return
	end if

	if (iter.ge.ITMAX)then
	write(*,*)'exceeds'
	stop !’ITMAX exceeded in amoeba’
	end if

	iter=iter+2

	ytry=amotry1(p,y,psum,mp,np,ndim,M,na,iu,iv,zp,zn,
     &  area1,ihi,-1.0d0)
	if (ytry.le.y(ilo)) then
	 ytry=amotry1(p,y,psum,mp,np,ndim,M,na,iu,iv,zp,zn,
     &   area1,ihi,2.0d0)
	else if (ytry.ge.y(inhi)) then
	 ysave=y(ihi)
	 ytry=amotry1(p,y,psum,mp,np,ndim,M,na,iu,iv,zp,zn,
     &   area1,ihi,0.5d0)
	 if (ytry.ge.ysave) then
	  do i=1,ndim+1
	   if(i.ne.ilo)then
	    do j=1,ndim
	     psum(j)=0.5d0*(p(i,j)+p(ilo,j))
	     p(i,j)=psum(j)
	    end do
	   y(i)=area1(M,na,iu,iv,zp,zn,psum)	
	   end if
	  end do
	  iter=iter+ndim
	  go to 1
	 end if
	else
	 iter=iter-1
	end if
	go to 2
	END

	FUNCTION amotry1(p,y,psum,mp,np,ndim,M,na,iu,iv,zp,zn,
     &  area1,ihi,fac)
	implicit double precision(a-h,o-z)
	dimension p(mp,np),psum(np),y(mp),na(M)
	PARAMETER (NMAX=20)
	EXTERNAL area1
	dimension ptry(NMAX)


	fac1=(1.0d0-fac)/ndim
	fac2=fac1-fac
	do j=1,ndim
	ptry(j)=psum(j)*fac1-p(ihi,j)*fac2
	end do
	ytry=area1(M,na,iu,iv,zp,zn,ptry) 	
	if (ytry.lt.y(ihi)) then
	y(ihi)=ytry
	do j=1,ndim
	psum(j)=psum(j)-p(ihi,j)+ptry(j)
	p(ihi,j)=ptry(j)
	end do
	end if
	amotry1=ytry	
	return
	END

	SUBROUTINE amoeba2(p,y,mp,np,ndim,ftol,M,na,iu,iv,zp,zn,
     &  area2,iter)
	implicit double precision(a-h,o-z)
	dimension p(mp,np),y(mp),na(M)
	PARAMETER (NMAX=20,ITMAX=5000,TINY=1.e-10) 
	EXTERNAL area2

	double precision psum(NMAX)	

	iter=0
1 	do n=1,ndim
	 sum=0. 
	 do m1=1,ndim+1
	  sum=sum+p(m1,n)
	 end do
	 psum(n)=sum
	end do

2 	ilo=1
	if (y(1).gt.y(2)) then 
	 ihi=1
	 inhi=2
	else
	 ihi=2
	 inhi=1
	end if

	do i=1,ndim+1
	 if(y(i).le.y(ilo)) ilo=i
	 if(y(i).gt.y(ihi)) then
	  inhi=ihi
	  ihi=i
	 else if(y(i).gt.y(inhi)) then
	 if(i.ne.ihi) inhi=i
	 end if
	end do


	rtol=2.*abs(y(ihi)-y(ilo))/(abs(y(ihi))+abs(y(ilo))+TINY)
c	write(*,*)rtol

	if (rtol.lt.ftol) then 
	 swap=y(1)
	 y(1)=y(ilo)
	 y(ilo)=swap
	 do n=1,ndim
	  swap=p(1,n)
	  p(1,n)=p(ilo,n)
	  p(ilo,n)=swap
	 end do
	 return
	end if

	if (iter.ge.ITMAX)then
	write(*,*)'exceeds'
	stop !’ITMAX exceeded in amoeba’
	end if

	iter=iter+2

	ytry=amotry2(p,y,psum,mp,np,ndim,M,na,iu,iv,zp,zn,
     &  area2,ihi,-1.0d0)
	if (ytry.le.y(ilo)) then
	 ytry=amotry2(p,y,psum,mp,np,ndim,M,na,iu,iv,zp,zn,
     &   area2,ihi,2.0d0)
	else if (ytry.ge.y(inhi)) then
	 ysave=y(ihi)
	 ytry=amotry2(p,y,psum,mp,np,ndim,M,na,iu,iv,zp,zn,
     &   area2,ihi,0.5d0)
	 if (ytry.ge.ysave) then
	  do i=1,ndim+1
	   if(i.ne.ilo)then
	    do j=1,ndim
	     psum(j)=0.5d0*(p(i,j)+p(ilo,j))
	     p(i,j)=psum(j)
	    end do
	   y(i)=area2(M,na,iu,iv,zp,zn,psum)	
	   end if
	  end do
	  iter=iter+ndim
	  go to 1
	 end if
	else
	 iter=iter-1
	end if
	go to 2
	END

	FUNCTION amotry2(p,y,psum,mp,np,ndim,M,na,iu,iv,zp,zn,
     &  area2,ihi,fac)
	implicit double precision(a-h,o-z)
	dimension p(mp,np),psum(np),y(mp),na(M)
	PARAMETER (NMAX=20)
	EXTERNAL area2
	dimension ptry(NMAX)

	fac1=(1.0d0-fac)/ndim
	fac2=fac1-fac
	do j=1,ndim
	ptry(j)=psum(j)*fac1-p(ihi,j)*fac2
	end do
	ytry=area2(M,na,iu,iv,zp,zn,ptry)
	if (ytry.lt.y(ihi)) then
	y(ihi)=ytry
	do j=1,ndim
	psum(j)=psum(j)-p(ihi,j)+ptry(j)
	p(ihi,j)=ptry(j)
	end do
	end if
	amotry2=ytry
	return
	END
	
	SUBROUTINE amoeba3(p,y,mp,np,ndim,ftol,M,na,iu,iv,zp,zn,
     &  area3,iter)
	implicit double precision(a-h,o-z)
	dimension p(mp,np),y(mp),na(M)
	PARAMETER (NMAX=20,ITMAX=5000,TINY=1.e-10) 
	EXTERNAL area3

	double precision psum(NMAX)	

	iter=0
1 	do n=1,ndim
	 sum=0. 
	 do m1=1,ndim+1
	  sum=sum+p(m1,n)
	 end do
	 psum(n)=sum
	end do

2 	ilo=1
	if (y(1).gt.y(2)) then 
	 ihi=1
	 inhi=2
	else
	 ihi=2
	 inhi=1
	end if

	do i=1,ndim+1
	 if(y(i).le.y(ilo)) ilo=i
	 if(y(i).gt.y(ihi)) then
	  inhi=ihi
	  ihi=i
	 else if(y(i).gt.y(inhi)) then
	 if(i.ne.ihi) inhi=i
	 end if
	end do


	rtol=2.*abs(y(ihi)-y(ilo))/(abs(y(ihi))+abs(y(ilo))+TINY)
c	write(*,*)rtol

	if (rtol.lt.ftol) then 
	 swap=y(1)
	 y(1)=y(ilo)
	 y(ilo)=swap
	 do n=1,ndim
	  swap=p(1,n)
	  p(1,n)=p(ilo,n)
	  p(ilo,n)=swap
	 end do
	 return
	end if

	if (iter.ge.ITMAX)then
	write(*,*)'exceeds'
	stop !’ITMAX exceeded in amoeba’
	end if

	iter=iter+2

	ytry=amotry3(p,y,psum,mp,np,ndim,M,na,iu,iv,zp,zn,	
     &  area3,ihi,-1.0d0)	
	if (ytry.le.y(ilo)) then
	 ytry=amotry3(p,y,psum,mp,np,ndim,M,na,iu,iv,zp,zn,	
     &   area3,ihi,2.0d0)	
	else if (ytry.ge.y(inhi)) then
	 ysave=y(ihi)
	 ytry=amotry3(p,y,psum,mp,np,ndim,M,na,iu,iv,zp,zn,	
     &   area3,ihi,0.5d0)	
	 if (ytry.ge.ysave) then
	  do i=1,ndim+1
	   if(i.ne.ilo)then
	    do j=1,ndim
	     psum(j)=0.5d0*(p(i,j)+p(ilo,j))
	     p(i,j)=psum(j)
	    end do
	   y(i)=area3(M,na,iu,iv,zp,zn,psum)	
	   end if
	  end do
	  iter=iter+ndim
	  go to 1
	 end if
	else
	 iter=iter-1
	end if
	go to 2
	END

	FUNCTION amotry3(p,y,psum,mp,np,ndim,M,na,iu,iv,zp,zn,	
     &  area3,ihi,fac)	
	implicit double precision(a-h,o-z)
	dimension p(mp,np),psum(np),y(mp),na(M)
	PARAMETER (NMAX=20)
	EXTERNAL area3	
	dimension ptry(NMAX)
	
	
	fac1=(1.0d0-fac)/ndim
	fac2=fac1-fac
	do j=1,ndim
	ptry(j)=psum(j)*fac1-p(ihi,j)*fac2
	end do
	ytry=area3(M,na,iu,iv,zp,zn,ptry) 	
	if (ytry.lt.y(ihi)) then
	y(ihi)=ytry
	do j=1,ndim
	psum(j)=psum(j)-p(ihi,j)+ptry(j)
	p(ihi,j)=ptry(j)
	end do
	end if
	amotry3=ytry	
	return
	END							
	
	SUBROUTINE amoeba4(p,y,mp,np,ndim,ftol,M,na,iu,iv,zp,zn,	
     &  area4,iter)	
	implicit double precision(a-h,o-z)
	dimension p(mp,np),y(mp),na(M)
	PARAMETER (NMAX=20,ITMAX=5000,TINY=1.e-10) 
	EXTERNAL area4	
	
	double precision psum(NMAX)	
	
	iter=0
1 	do n=1,ndim
	 sum=0. 
	 do m1=1,ndim+1
	  sum=sum+p(m1,n)
	 end do
	 psum(n)=sum
	end do
	
c	write(*,*)'hello amoeba 1'	

2 	ilo=1
	if (y(1).gt.y(2)) then 
	 ihi=1
	 inhi=2
	else
	 ihi=2
	 inhi=1
	end if

	do i=1,ndim+1
	 if(y(i).le.y(ilo)) ilo=i
	 if(y(i).gt.y(ihi)) then
	  inhi=ihi
	  ihi=i
	 else if(y(i).gt.y(inhi)) then
	 if(i.ne.ihi) inhi=i
	 end if
	end do

c	write(*,*)'hello amoeba 2'		

	rtol=2.*abs(y(ihi)-y(ilo))/(abs(y(ihi))+abs(y(ilo))+TINY)
c	write(*,*)rtol

	if (rtol.lt.ftol) then 
	 swap=y(1)
	 y(1)=y(ilo)
	 y(ilo)=swap
	 do n=1,ndim
	  swap=p(1,n)
	  p(1,n)=p(ilo,n)
	  p(ilo,n)=swap
	 end do
	 return
	end if

	if (iter.ge.ITMAX)then
	write(*,*)'exceeds'
	stop !’ITMAX exceeded in amoeba’
	end if

	iter=iter+2

	ytry=amotry4(p,y,psum,mp,np,ndim,M,na,iu,iv,zp,zn,	
     &  area4,ihi,-1.0d0)	
	if (ytry.le.y(ilo)) then
	 ytry=amotry4(p,y,psum,mp,np,ndim,M,na,iu,iv,zp,zn,	
     &   area4,ihi,2.0d0)	
	else if (ytry.ge.y(inhi)) then
	 ysave=y(ihi)
	 ytry=amotry4(p,y,psum,mp,np,ndim,M,na,iu,iv,zp,zn,	
     &   area4,ihi,0.5d0)	
	 if (ytry.ge.ysave) then
	  do i=1,ndim+1
	   if(i.ne.ilo)then
	    do j=1,ndim
	     psum(j)=0.5d0*(p(i,j)+p(ilo,j))
	     p(i,j)=psum(j)
	    end do
	   y(i)=area4(M,na,iu,iv,zp,zn,psum)	
	   end if
	  end do
	  iter=iter+ndim
	  go to 1
	 end if
	else
	 iter=iter-1
	end if
	go to 2
	END

	FUNCTION amotry4(p,y,psum,mp,np,ndim,M,na,iu,iv,zp,zn,	
     &  area4,ihi,fac)	
	implicit double precision(a-h,o-z)
	dimension p(mp,np),psum(np),y(mp),na(M)
	PARAMETER (NMAX=20)
	EXTERNAL area4	
	dimension ptry(NMAX)

c	write(*,*)'hello amotry'	
	
	fac1=(1.0d0-fac)/ndim
	fac2=fac1-fac
	do j=1,ndim
	ptry(j)=psum(j)*fac1-p(ihi,j)*fac2
	end do
	ytry=area4(M,na,iu,iv,zp,zn,ptry) 	
	if (ytry.lt.y(ihi)) then
	y(ihi)=ytry
	do j=1,ndim
	psum(j)=psum(j)-p(ihi,j)+ptry(j)
	p(ihi,j)=ptry(j)
	end do
	end if
	amotry4=ytry	
	return
	END		
	
	include 'ran2.f'	
```
